# Synchronization properties in *C. elegans*: Relating behavioral circuits to structural and functional neuronal connectivity

**DOI:** 10.64898/2026.03.23.713580

**Authors:** Gourab Kumar Sar, Andrew Patton, Emma Towlson, Jörn Davidsen

## Abstract

A central question in neuroscience is how neural processing generates or encodes behavior. *Caenorhabditis elegans* is well suited to addressing this question, given its compact nervous system and near-complete structural connectome. Despite this, findings from previous studies remain inconclusive. While some have shown that the connectome can robustly encode specific behaviors such as locomotion, others report that functional connectivity can be reconfigured across behaviors. We aim to understand the relationship between structural connectivity, functional connectivity and biological behavior *in silico* by using an experimentally motivated computational model leveraging the structural connectome. Stimulation of specific neurons in the model induces oscillatory neural responses, enabling us to infer neuronal functional connectivity. Functional connectivity is found to be stronger among some neurons, allowing us to identify functional communities. We find that electrical synapses play a critical role in determining functional communities, and the resulting mesoscale functional architecture is predominantly gap junctionally assortative. Furthermore, comparison with behavioral circuits shows that locomotion circuits are largely segregated into distinct functional communities while other circuits are more distributed across multiple functional communities. We also observe that stimulation of neurons belonging to these distributed circuits elicits a more synchronized neuronal response compared to stimulation of neurons within the more segregated circuits. This is consistent with the presence of behavioral patterns that originate in one circuit and terminate in another (e.g., chemosensation leading to locomotion), such that stimulation of one circuit can activate the other and eventually result in a synchronized response. We also find a large repertoire of chimera-like synchronization patterns upon stimulation of certain behavioral circuits (chemosensation, mechanosensation) indicating high dynamical flexibility. Overall, our results demonstrate that while certain behaviors are governed by functionally segregated circuits, others emerge from the synchronization of multiple functional communities, which are, to begin with, influenced by the underlying structural connectivity.

**Author summary:** Animals constantly transform sensory inputs into actions, but it is still unclear how this mapping from neural activity to behavior is implemented in a real nervous system. *Caenorhabditis elegans* offers a unique testbed for this question because its entire wiring diagram is nearly completely mapped. Yet, previous works have reached mixed conclusions about how well this anatomical circuit diagram predicts actual patterns of activity and behavior. Here, we use a biologically inspired computational model of the *C. elegans* nervous system to bridge this gap between structure, function, and behavior. By virtually stimulating individual neurons and observing the resulting network-wide oscillations, we infer how strongly different pairs and groups of neurons interact in functional terms. We then use network analysis tools to identify groups of neurons that tend to co-activate, and relate these functional communities to known behavioral circuits for locomotion and sensory processing. We find that gap junctions play a key role in shaping functional communities, and that locomotion-related neurons are more functionally segregated than neurons involved in other behaviors, which are more functionally distributed. Our results suggest that some behaviors rely on specialized, functionally isolated circuits, whereas others emerge from the coordinated activity of multiple functional communities.

## Introduction

Understanding how neural activity gives rise to behavior is a central problem in neuroscience. This question is especially tractable in the nematode *Caenorhabditis elegans*, whose nervous system consists of a total of 302 neurons and for which an almost complete structural connectome is available [1–6]. Despite this exceptional level of anatomical detail, the exact relationship between structural connectivity, functional dynamics, and behavior in *C. elegans* remains unresolved. Several complementary approaches have been used to probe the relationship between structure, function, and behavior, ranging from purely *in silico* connectome-based modeling [7–9] and experimental measurements of neural activity and perturbations [10–12], to hybrid methods that explicitly integrate computational models with experimental data [13, 14].

Notably, specific structural features can robustly predict functional organization and behaviorally relevant computations, particularly in locomotor circuitry. On the experimental side, brain-wide calcium imaging has shown that graph-theoretic features of the anatomical connectome, especially hub neurons and non-local connectivity, are strongly associated with global functional correlations, and that perturbing/inhibiting multiple hub neurons selectively disrupts this brain-wide correlation structure [12].

Symmetry-based analyses of the connectome further identify structured “equivalence classes” (symmetry sectors) that constrain coordinated dynamics and yield functionally meaningful partitions in locomotion circuits [15]. In the same spirit, another work that explicitly couples connectome structure to whole-neural activity argues that network symmetries (e.g., fibration symmetries) help explain which subsets of neurons synchronize during locomotor control [16]. Complementing these graph-theoretic perspectives, dynamical and control-theoretic studies show that the connectome can encode low-dimensional motor programs and can predict neurons critical for locomotion, linking structural controllability or feedback motifs to experimentally testable behavioral outcomes [17–19].

Behavior [20–22] provides a natural bridge between anatomical structure and functional dynamics: while the connectome supplies a relatively stable scaffold, functional coupling is flexible and can be reconfigured across behavioral contexts, internal states, and timescales [23, 24]. Whole-brain calcium imaging shows that a large fraction of the nervous system participates in coordinated, low-dimensional population dynamics that track locomotor commands and motor-state transitions, linking brain-wide functional organization to a defined behavioral repertoire [10]. Likewise, imaging in freely moving *C. elegans* reveals reproducible activity patterns underlying persistent foraging states such as roaming versus dwelling, illustrating state-dependent reorganization of functional connectivity on the same structural substrate [24]. Taken together, prior works have yielded a diverse set of results linking structure, function, and behavior. Reconciling these complementary, and sometimes divergent, perspectives requires a framework that explicitly connects anatomical wiring, neuronal dynamics, and behavioral output.

A natural way to build such a unifying framework is through computational models that translate anatomical wiring into whole-brain neuronal dynamics. In this spirit, a wide range of computational approaches has been used to study whole-brain neuronal dynamics in *C. elegans* [25]. Earlier modeling studies often employed spiking neuron models, such as the Hindmarsh-Rose system, to investigate synchronization phenomena in the connectome [26, 27]. However, this modeling choice is biologically questionable, as neurons in *C. elegans* and other nematodes largely lack classical action potentials and instead communicate via graded membrane potential changes [28, 29]. More recent work has therefore shifted toward biophysically motivated membrane models that better capture the experimentally observed voltage dynamics of *C. elegans* neurons [17, 30, 31]. Parallel to developments in neuronal modeling, several network theoretic approaches have been applied to relate structure and function in the *C. elegans* connectome. These include symmetry-based analyses [15, 32], modularity maximization methods such as Louvain community detection [33], and stochastic block modeling frameworks [11, 26]. Collectively, these studies suggest that mesoscale organization (e.g., modules or communities, hubs, symmetry-defined neuron groups) strongly shapes the emergent neural dynamics by constraining how activity co-fluctuates and propagates across the network, rather than being determined solely by single connections. Nevertheless, there is no consensus on whether and how structural modules correspond to functional units or behavioral circuits.

In this study, we aim to clarify how structural connectivity shapes functional organization and synchronization patterns in the *C. elegans* nervous system, and how these functional structures relate to well-characterized behavioral circuits. To this end, we employ a biologically motivated single-compartment membrane model that explicitly incorporates both chemical synapses and electrical gap junctions as given by the anatomical connectome [30]. By systematically stimulating individual neurons *in silico*, we induce network-wide oscillatory responses and use these responses to infer functional connectivity from pairwise correlations of membrane potentials. Rather than focusing solely on individual neuron pairs, we investigate the mesoscale functional architecture of the network. Using a weighted stochastic block model, we identify functional communities – groups of neurons that respond similarly to stimulation – across an ensemble of stimulation conditions. This approach allows us to determine how functional communities emerge from the static connectome and to assess the relative structural roles in shaping these communities. We then relate the detected functional communities to five well-studied behavioral circuits: forward and backward locomotion [34–36], chemosensation [37–40], mechanosensation [41–44], and klinotaxis [45–47]. These circuits differ markedly in their anatomical composition and behavioral roles. Locomotion circuits are organized to generate robust and repeatable motor programs, while sensory circuits – chemosensory, mechanosensory, and klinotaxis pathways – are organized to flexibly transform rich external cues into state and context-dependent neural responses [21, 34, 48]. By comparing how these circuits map onto functional communities and by analyzing their synchronization patterns under targeted stimulation, we assess whether different behavioral modalities rely on functionally segregated or more distributed neural architectures.

Our results show that functional communities in the *C. elegans* nervous system are strongly shaped by electrical synapses (or gap junctions) and are predominantly gap junctionally assortative. Locomotion circuits are largely confined to specific functional communities, consistent with their specialized role, whereas the sensory circuits are distributed across multiple communities. We find that stimulation to these distributed sensory circuits leads to stronger global synchronization than stimulation of more segregated locomotion circuits. Moreover, we observe a broad repertoire of response patterns upon stimulating some of these behavioral circuits, characterized by the coexistence of in-phase and out-of-phase responses. The large number of such chimera patterns [49–51] signifies a large degree of dynamical flexibility of these behavioral circuits. Together, our findings provide a unified picture of how structure, function, and behavior might be linked in *C. elegans* and highlight the central role of electrical coupling in organizing functional brain architecture.

## Materials and methods

### Structural connectome dataset

The *C. elegans* nervous system is one of the very few fully-mapped connectomes of any organism. An adult hermaphrodite nervous system consists of a total of 302 neurons, of which 279 are somatic neurons, 20 are pharyngeal neurons, and 3 neurons do not have any synaptic connection to others. The somatic nervous system is anatomically connected through electrical (gap junction) and chemical synapses. Gap junctions form undirected connections, whereas chemical synapses are directed. In total, the connectome comprises 6,393 chemical synapses and 890 gap junctions. The neuronal network is weighted, as some neuron pairs are connected by multiple gap junctions or synapses. There are 2,990 unique weighted synaptic connections when chemical synapses and gap junctions are considered together. For further details about the structural connectome, we refer the reader to Refs. [1–3, 52].

Although there is still a lack of electrophysiological experimental evidence confirming whether chemical synapses between neurons are excitatory or inhibitory [53], it is commonly assumed that there are 26 GABAergic neurons that contribute to inhibitory synapses. The remaining neurons (primarily glutamatergic and cholinergic) are generally considered excitatory [2]. Neurons are broadly classified into three categories based on their structural properties and functional roles: sensory neurons, interneurons (or command neurons), and motor neurons. Sensory neurons respond to specific environmental stimuli and convert them into neural signals; interneurons integrate and process this information, coordinating the appropriate neural response; and motor neurons relay signals to muscles, ultimately generating the organism’s behavioral output.

Some neurons exhibit multiple functional roles and are therefore classified as polymodal neurons. The somatic nervous system of *C. elegans* consists of 68 sensory neurons, 67 interneurons, 89 motor neurons, and 55 polymodal neurons. See Fig 1 for a schematic of the *C. elegans* neuronal network where the neurons are colored according to their respective types.

**Fig 1.**
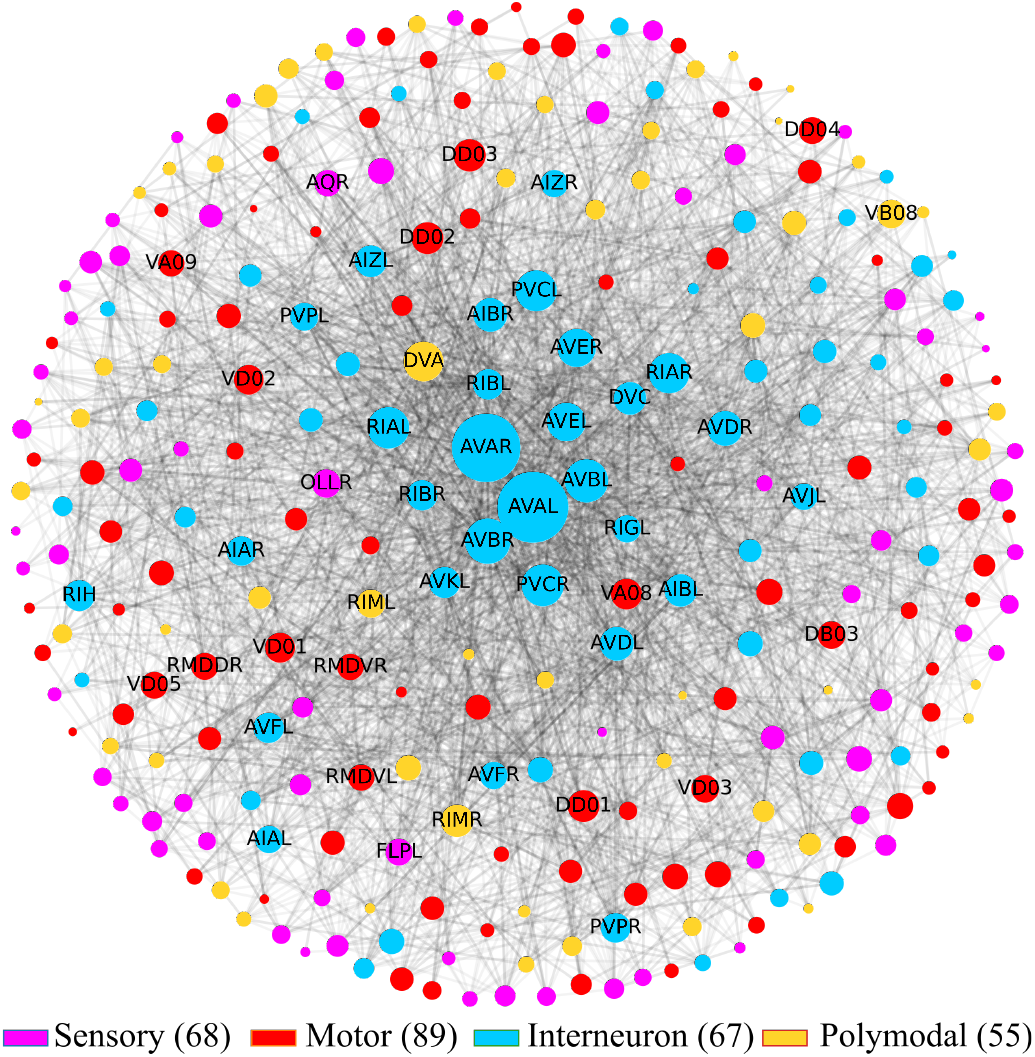
*C. elegans* neuronal network consisting of 279 somatic neurons. The edges represent the gap junctions and chemical synapses. Node sizes in the structural connectome are scaled according to total degree, and nodes are colored according to their respective types. Labels are shown for the 50 neurons with the highest degrees.

### Behavioral circuits

*C. elegans* exhibits a broad spectrum of behaviors, ranging from locomotion, foraging, feeding, and touch withdrawal to sensory-driven taxis based on odor, taste, and temperature [21, 22]. A diverse toolkit is available for investigating and manipulating these behaviors, including automated visual tracking, microfluidic platforms that finely control structured environments, optogenetics, calcium imaging, laser ablation, and electrophysiological and genetic methods that enable precise interrogation of neural circuits [54–58]. In the following, we briefly describe several key behavioral modalities through which *C. elegans* interacts with its environment and discuss the neuronal circuits associated with each of these behaviors. Table 1 provides the full list of neurons in the different behavioral circuits – they share some of their constituent neurons (Table A in S1 Appendix) – and Fig 2 shows the proportions of the different types of neurons the circuits contain.

**Table 1.** Neuronal composition of five behavioral circuits.

| Circuit | Neurons |
| --- | --- |
| Forward locomotion | AVBL, AVBR, DB1, DB2, DB3, DB4, DB5, DB6, DB7, PVCL, PVCR, RIBL, RIBR, VB1, VB2, VB3, VB4, VB5, VB6, VB7, VB8, VB9, VB10, VB11 |
| Backward locomotion | AIBL, AIBR, AVAL, AVAR, AVDL, AVDR, AVEL, AVER, DA1, DA2, DA3, DA4, DA5, DA6, DA7, DA8, DA9, RIML, RIMR, VA1, VA2, VA3, VA4, VA5, VA6, VA7, VA8, VA9, VA10, VA11, VA12 |
| Chemosensation | ADFL, ADFR, ADLL, ADLR, AIAL, AIAR, AIBL, AIBR, AIYL, AIYR, AIZL, AIZR, ASEL, ASER, ASGL, ASGR, ASHL, ASHR, ASIL, ASIR, ASJL, ASJR, ASKL, ASKR, AVAL, AVAR, AVBL, AVBR, AVDL, AVDR, AVEL, AVER, AWAL, AWAR, AWBL, AWBR, AWCL, AWCR, PHAL, PHAR, PHBL, PHBR, PVCL, PVCR, RIML, RIMR |
| Mechanosensation | ADEL, ADER, ALML, ALMR, ASHL, ASHR, AVAL, AVAR, AVBL, AVBR, AVDL, AVDR, AVM, CEPDL, CEPDR, CEPVL, CEPVR, FLPL, FLPR, IL1DL, IL1DR, IL1L, IL1R, IL1VL, IL1VR, OLQDL, OLQDR, OLQVL, OLQVR, PDEL, PDER, PLML, PLMR, PVCL, PVCR, PVDL, PVDR, PVM |
| Klinotaxis | ADFR, AIAL, AIAR, AIBL, AIBR, AIML, AIYL, AIYR, AIZL, AIZR, ASEL, ASER, AWAR, AWBR, PVT, RIBL, RIBR, RMGL, SAADL, SMBDL, SMBDR, SMBVL, SMBVR |

**Fig 2.**
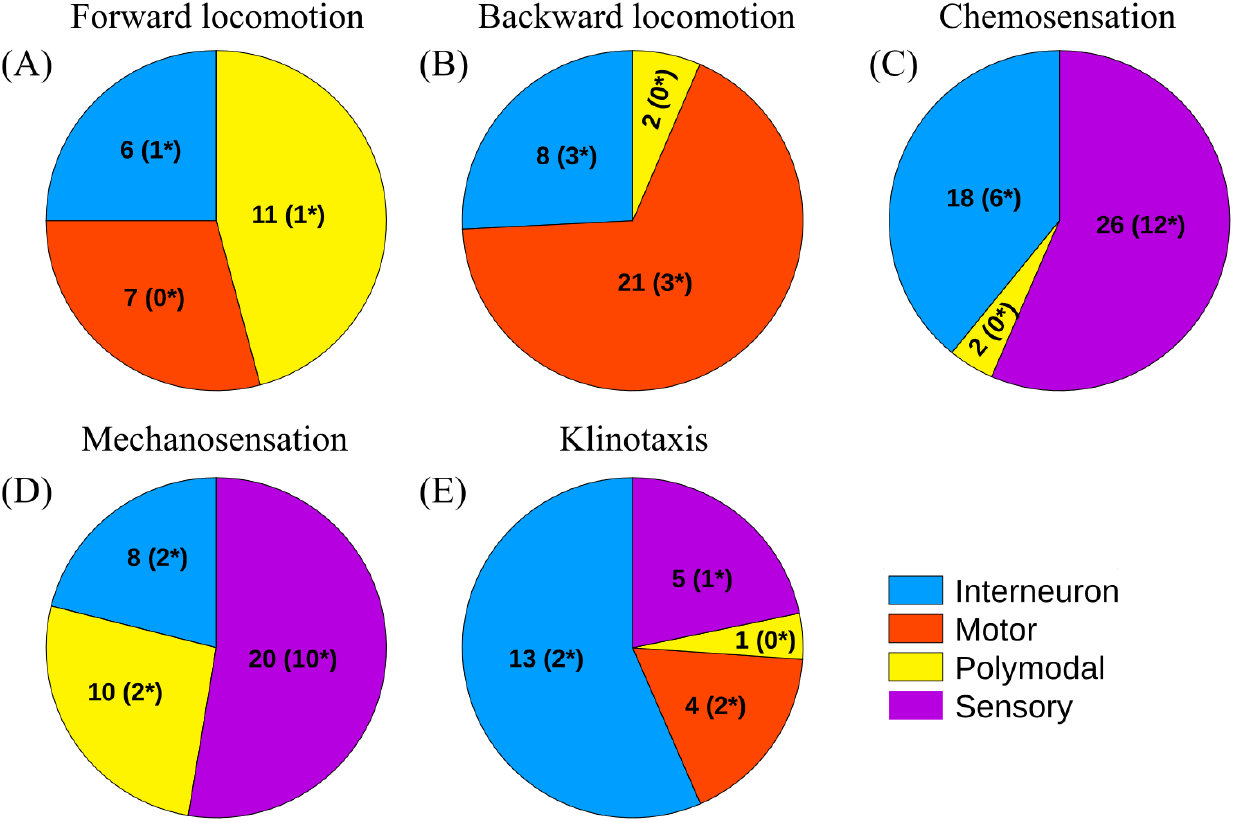
Composition of behavioral circuits. The proportions of interneurons, sensory neurons, motor neurons, and polymodal neurons are shown in (A) forward locomotion, (B) backward locomotion, (C) chemosensation, (D) mechanosensation, and (E) klinotaxis circuits. Inside the parenthesis, the number of *susceptible* neurons (neurons that facilitate oscillatory responses from other neurons upon stimulation in our mathematical model) are shown. In total, these five circuits are made up of 123 unique neurons, with some of them being present in multiple communities. Of these 123 neurons, 34 neurons are susceptible neurons with the majority being sensory neurons (19 out of 34).

### Locomotion

*C. elegans* moves primarily by generating sinusoidal waves along its body, producing forward movement under normal conditions. Reversals, followed by reorientation maneuvers such as omega turns, allow the worm to change direction and escape adverse stimuli. Distinct motor circuits control forward and backward motion: forward locomotion is primarily driven by the interneurons AVB and PVC, whereas backward movement is initiated by the interneurons AVA, AVD, and AVE. These command interneurons coordinate with ventral and dorsal motor neurons to enable rapid and flexible transitions between locomotor states [34–36]. As Fig 2 shows, whereas the locomotion circuits comprise a large number of motor neurons, the other circuits contain a larger proportion of sensory neurons or interneurons – consistent with the nature of the tasks they perform.

### Chemosensation

Chemosensation enables *C. elegans* to detect and navigate chemical gradients associated with food, pheromones, and harmful substances [37–40]. This behavior is mediated primarily by the amphid sensory neurons, which include several well-characterized chemosensory cell types. The neurons AWA and AWC detect attractive odorants, while ASE senses water-soluble attractants such as salts and amino acids. Avoidance of repellents is mediated largely by ASH, ADF, and ASK, which respond to noxious chemicals and high osmolarity. These sensory inputs are transmitted to downstream interneurons, including AIA, AIB, and AIY, which integrate chemical information and drive appropriate motor outputs for chemotaxis. Together, this network enables the worm to perform robust attraction, repulsion, and navigation in complex chemical environments.

### Mechanosensation

*C. elegans* relies on a well-characterized mechanosensory system to detect gentle and harsh touch, enabling rapid protective and exploratory responses [41–44]. Gentle touch along the body is sensed by a set of six touch receptor neurons - ALML/R and AVM in the anterior, and PLML/R and PVM in the posterior - which mediate direction-specific withdrawal or acceleration. These neurons use specialized MEC-4/MEC-10 mechanotransduction channels to convert mechanical stimuli into electrical signals.

Harsh or noxious mechanical stimuli are detected by additional polymodal neurons, including PVD and FLP, which elicit stronger escape responses. Together, these mechanosensory pathways provide the worm with a robust ability to respond to physical perturbations and navigate complex environments.

### Klinotaxis

Klinotaxis in *C. elegans* refers to a navigation strategy in which the worm steers toward or away from a stimulus by comparing changes in concentration over successive head swings rather than sampling spatial differences across its body [45–47]. In chemical gradients, the worm adjusts the curvature of its trajectory based on temporal changes in stimulus intensity sensed during these head sweeps. A well-studied example is salt (NaCl) klinotaxis, where the bilateral ASE chemosensory neurons (ASEL/ASER) detect changes in NaCl concentration and relay this information to downstream interneurons such as AIY and AIZ, which in turn drive neck motor neurons of the SMB class to bias head bending toward preferred salt concentrations.

### Single-compartment membrane model of *C. elegans*

In our model approach, each neuron in the *C. elegans* somatic nervous system is modeled as a single electrical compartment whose membrane potential evolves continuously in response to synaptic and gap junction inputs following [30]. This model is inspired by *in-situ* membrane voltage recordings, which show that neuronal responses are graded potentials, making them more appropriate for capturing voltage dynamics than conventional multi-compartment spiking neuron models. The model incorporates the biophysical properties of chemical synapses and electrical couplings through weighted anatomical connectivity. This framework allows us to simulate how external stimuli propagate through the network and to examine the emergence of collective neuronal responses across the full connectome. The nonlinear dynamics of neuronal activity is governed by a pair of equations [30]:

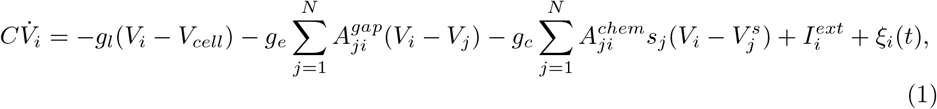

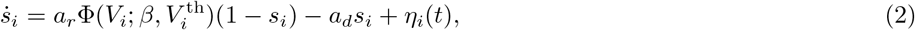

where *V*_*i*_ is the membrane potential of the *i*-th neuron. *C* denotes the membrane capacitance, *g*_*l*_ the membrane leakage conductance, and *V*_*cell*_ the leakage potential. External stimulation contributes through the current 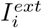, whereas interactions within the network arise from gap junction currents and synaptic currents. Gap junctions are modeled as ohmic connections, with 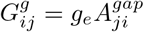 denoting the total conductance between neurons *i* and *j*. Chemical synapses are described using the maximal synaptic conductance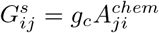, scaled by the synaptic activity variable *s*_*i*_, and their contribution depends on the deviation of the postsynaptic voltage from the corresponding synaptic reversal potential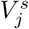, which is a constant but with different values for excitatory and inhibitory presynaptic neurons. Here, 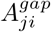and 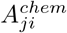are the weighted gap junction and chemical synapse connectivity, respectively, obtained directly from the structural connectome dataset described above. The evolution of synaptic activity is governed by rise and decay times *a*_*r*_ and *a*_*d*_, together with a sigmoid activation function of width *β*. The activation function Φis given by

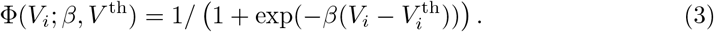

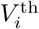is the threshold voltage for each neuron which is calculated by setting Φ= 1*/*2 at equilibrium where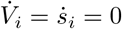. Typically, this threshold potential is determined by setting 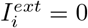[59]. Here, however, we calculate the threshold using a nonzero external current, so that 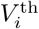becomes a function of 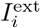(similar to Refs. [30, 32]).

We also include Gaussian white noise terms, *ξ*_*i*_(*t*) and *η*_*i*_(*t*), in the equations, with noise strength *σ*. In our simulations, we set *σ* = 5 × 10^*−*5^. Table 2 lists the parameters used in the above system, following the standard values reported in Ref. [59].

**Table 2.** Neuronal system parameters.

| Parameters | Descriptions | Numerical values with units |
| --- | --- | --- |
| $C$ | Whole-cell membrane capacitance | 1pF |
| $V_{cell}$ | Leakage potential | -35 mV |
| $g_l$ | Leakage conductance | 10 pS |
| $g_e$ | Gap junction conductance | 100 pS |
| $g_c$ | Chemical synapse conductance | 100 pS |
| $\alpha_l = g_l/C$ | Membrane potential time constant | 10 s<sup>-1</sup> |
| $\alpha_e = g_e/C$ | Gap junction time constant | 100 s<sup>-1</sup> |
| $\alpha_c = g_c/C$ | Chemical synapse time constant | 100 s<sup>-1</sup> |
| $V^s$ | Reversal potentials | $\begin{cases} 0 \text{ mv for excitatory presynaptic neurons,} \\ -45 \text{ mv for inhibitory presynaptic neurons.} \end{cases}$ |
| $a_r$ | Synaptic activity rise time | 1 s<sup>-1</sup> |
| $a_d$ | Synaptic activity decay time | 5 s<sup>-1</sup> |
| $\beta$ | Steepness of the sigmoid ($\Phi$) function | 0.125 mV<sup>-1</sup> |
| $I_i^{ext}$ | External input current | variable (pA) |

A complete list of parameters and their descriptions for the single-compartment membrane model.

#### Numerics

For numerical simulations, we employ the Euler-Maruyama integration method with a time step of *dt* = 10^*−*4^. The initial conditions are set by perturbing *V*_*i*_ and *s*_*i*_ from their equilibrium values by a small amplitude *ϵ* = 2.5× 10^*−*3^. We have also verified the robustness of the results with respect to the choice of initial conditions by using nonidentical perturbations drawn from a Gaussian distribution centered around 0 with a standard deviation of 2.5 × 10^*−*3^. We ran simulations up to *T* = 200 s and analyzed only the last 20% of the data, which ensured that transients had settled and that each neuron exhibited a sufficient number of oscillations if present.

### Emergent dynamics

Previous studies have reported that the fixed point of the system is globally stable in the absence of external input currents [17, 31]. Moreover, a constant, explicit and sufficiently large external input is required for the system to enter an oscillatory state and sustain those oscillations (i.e., oscillations that do not decay over time). Therefore, we apply a constant stimulus 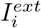on the *i*-th neuron and observe the long-term dynamics of the system. If the stimulus is too large, the oscillations become chaotic in some cases. Excluding these large stimuli, we observe that one of the following three scenarios occurs:

1. The equilibrium remains stable.
2. The equilibrium becomes unstable and the trajectory converges to a new fixed point.
3. The trajectory converges to a stable limit cycle via a supercritical Hopf bifurcation; see Fig 3B for an example.

**Fig 3.**
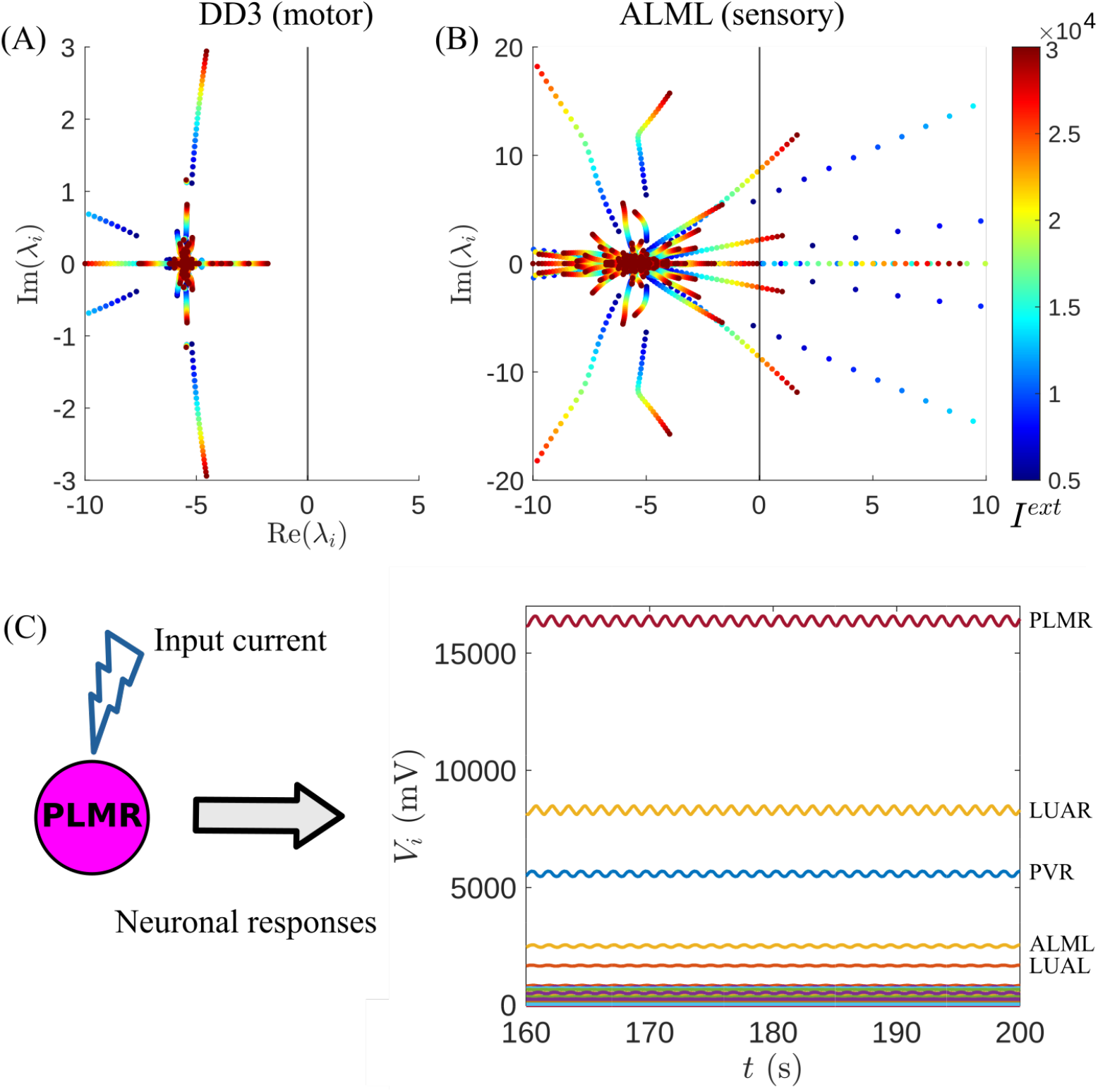
Spectrum of eigenvalues as a function of input current amplitude. (A) The motor neuron DD03 is stimulated and it is found that the equilibrium is always stable. (B) The sensory neuron ALML is stimulated and the system goes to an oscillatory state via a supercritical Hopf bifurcation. The *x*-axis, which denotes the real parts of the eigenvalues, is truncated for visual clarity. (C) Example of the voltage responses of all neurons when a single neuron (here the PLMR neuron) is stimulated with an external current in our computational model. The noise strength *σ* is set to zero here.

The linear stability of the noise-free system is analyzed by evaluating the eigenvalues of the Jacobian matrix at the fixed point. In Fig 3A-B, the eigenvalue spectra are shown as functions of the input current amplitude for two different cases. It is evident that both the input current magnitude and the stimulated neuron are crucial in determining which of the three scenarios described above will be encountered. For example, consider stimulation with an external input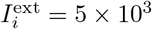. Under this stimulus, the system remains at equilibrium in 95 out of 279 cases, converges to a new fixed point in 161 cases, and enters an oscillatory state in 23 cases. When the stimulus is increased to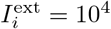, these numbers change to 46 (equilibrium), 203 (new fixed point), and 30 (oscillatory). We also find that for sufficiently large input currents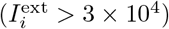, the oscillations become chaotic in some cases. To avoid this regime, we restrict 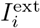 to a range in which only periodic oscillatory behavior and stable equilibria are observed.

In the oscillatory state, all neurons (except 4 neurons – IL2DL, IL2DR, PLNR, PVDR – which have neither gap junctions nor inward synaptic connections with the rest of the network) begin to oscillate with identical frequencies. This holds for both variables *V*_*i*_ and *s*_*i*_. Although the neurons are frequency-locked, their oscillations differ significantly in amplitude and range (see Fig 3C, which shows the voltage responses following PLMR stimulation). In Fig. A of S1 Appendix, we show the range of the response amplitudes for both variables under stimulation with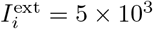. For a more detailed discussion of the nonlinear dynamics of the system, we refer the reader to Refs. [17, 31].

### Phase calculation and synchronization measure

Since the membrane potentials of neurons and the associated synaptic activity variables vary in both range and amplitude, we normalize them to the interval [−1, 1] by subtracting the mid-range value (to center the signal) and dividing by the half-range (to scale the amplitude) so that the oscillations remain within a comparable scale. We then define the phase of the *i*-th neuron as [49, 50]

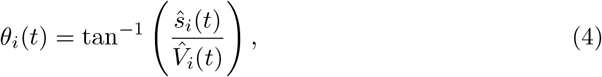

where 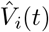 and *ŝ*_*i*_(*t*) are the normalized variables (the MATLAB function atan2 is used for this purpose). Once we have a phase description of the neurons, we can examine whether they tend to be in-phase or out-of-phase. This is especially useful because the neurons are frequency-locked, and it is their phases that separate them or make them heterogeneous. To quantify the global phase synchronization, we use the Kuramoto order parameter, defined as

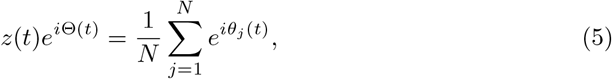

and consider its time average *Z* = ⟨*z*(*t*)⟩ _*T*_. By definition, *Z* ranges from 0 to 1, where 0 corresponds to a state where the phases are uniformly distributed and 1 corresponds to a fully in-phase state.

However, it is also observed that certain groups of neurons behave more similarly (or differently) in their phases compared to other groups of neurons. This necessitates capturing these group-wise relationships. We accomplish this by defining a Kuramoto order parameter between the *k*-th and *l*-th groups as

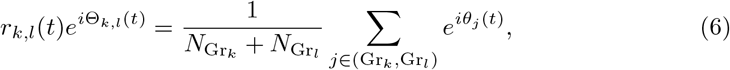

where Gr_*i*_ and 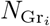 denote the *i*-th group and the number of neurons it contains, respectively. The time averaged Kuramoto order parameter is then calculated as

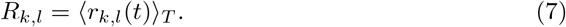

Like *Z, R* also ranges from 0 to 1, with larger values corresponding to a higher level of synchrony or a phase-coherent state. In Fig 4A, we show the Kuramoto order matrix when the AVEL neuron in the backward locomotion circuit is stimulated. Here, the groups correspond to the behavioral circuits, and it is observed that the synchronization between the behavioral circuits is relatively low. In Fig 4B, we present the example time series of the phase difference with the AVEL neuron for different neurons from different behavioral circuits. The aforementioned lower level of synchronization is visible in these time series.

**Fig 4.**
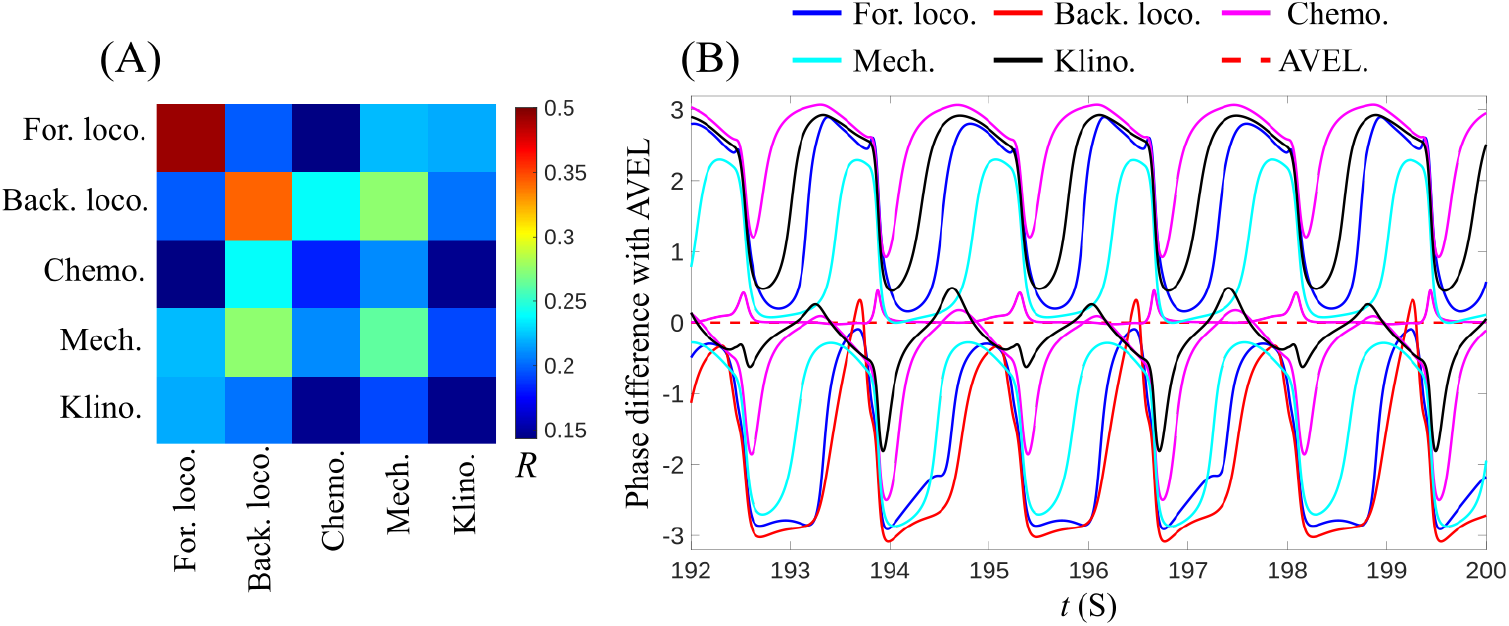
Kuramoto order matrix and phase difference time series. (A) Kuramoto order matrix *R*_*k,l*_ for the behavioral circuits when the AVEL neuron (interneuron) in the backward locomotion circuit is stimulated. The colorbar indicates the value of the matrix entry. (B) Time series of the phase differences with the stimulated AVEL neuron. Two example neurons from each behavioral circuit are shown: blue – forward locomotion, red – backward locomotion, magenta – chemosensation, cyan – mechanosensation, and black – klinotaxis.

### Correlation based functional community detection

To evaluate functional connectivity in our model, we compute the pairwise Pearson correlation between the voltage responses when stimulating a single neuron. Let *P* denote the correlation matrix, where *P*_*ij*_ is the Pearson correlation between *V*_*i*_(*t*) and *V*_*j*_(*t*), given by

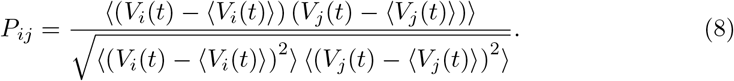

We then apply the weighted stochastic block model (WSBM) [60, 61] to detect communities from the correlation matrix *P*. For this, we use the MATLAB implementation of WSBM developed by A. Clauset [62], which allows the number of communities to be specified. The left panel of Fig 5 illustrates the WSBM approach, showing both the original correlation matrix and the WSBM-sorted matrix when the number of communities is fixed to six.

**Fig 5.**
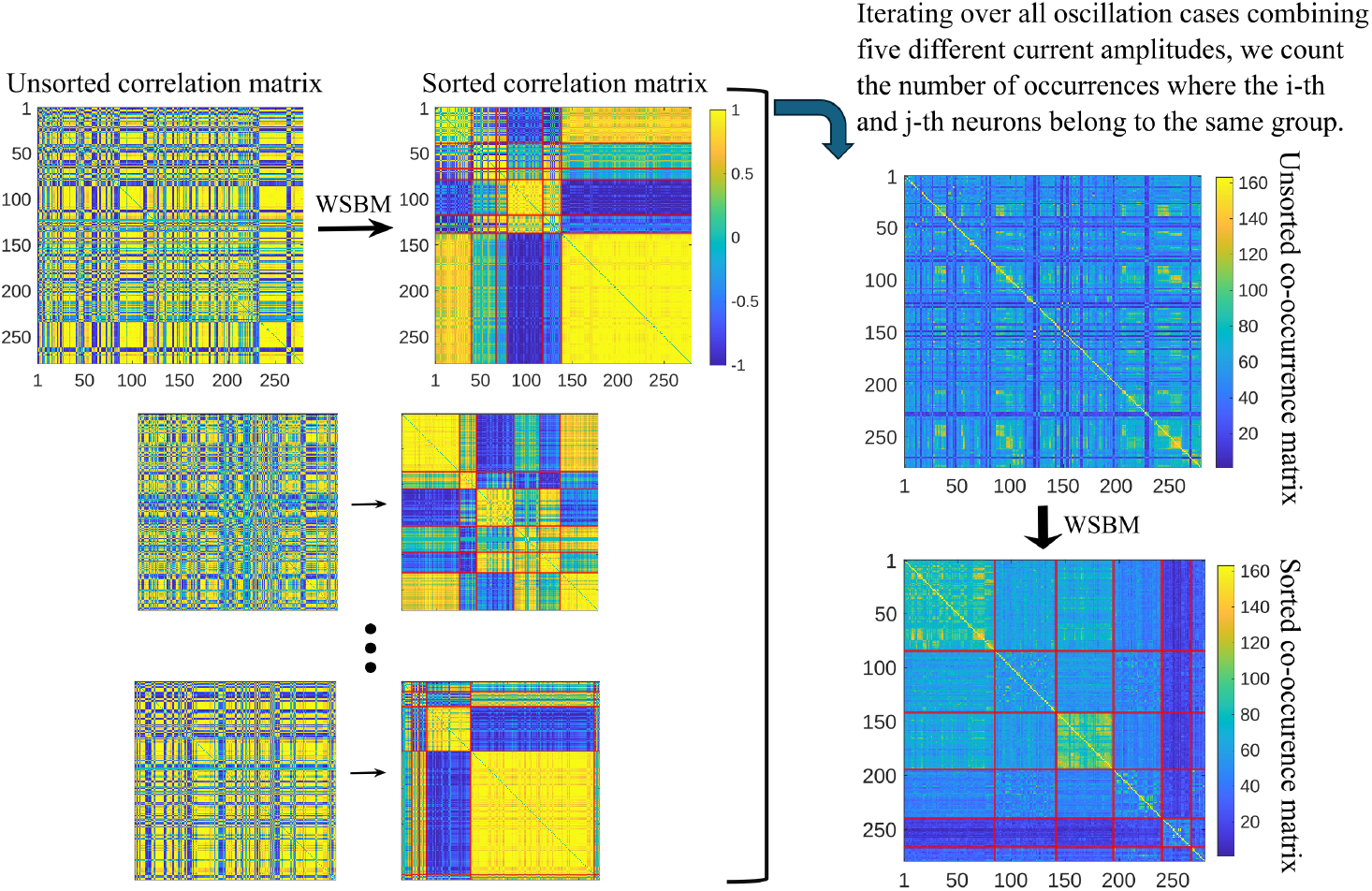
Functional community detection using the WSBM method. Left panel: WSBM-based community detection applied to 163 Pearson correlation matrices. Neurons are assigned to six communities in each instance. Right panel: The number of times any two neurons appear in the same community is counted to form the co-occurrence matrix *Q*. WSBM is then applied to *Q* to obtain the final six functional communities.

Since the correlation matrix and, hence, the detected communities depend on which neuron is stimulated and how strongly it is stimulated, we pursue an ensemble approach similar to Ref. [26]. Neurons are stimulated with five different current amplitudes, and we collect all instances in which oscillatory responses occur. In total, we obtain 163 instances of oscillatory neuronal responses (23 for 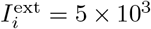, 26 for 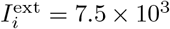, 30 for 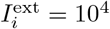, 44 for 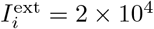, and 40 for 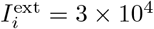) resulting in 163 corresponding *P* matrices that include 78 unique neurons. We apply WSBM to each of these matrices, detecting the same number of communities each time. We then count the number of occurrences in which any two neurons appear in the same community, yielding a co-occurrence matrix *Q* with entries ranging from 0 to 163. Finally, we apply WSBM again to this co-occurrence matrix *Q* to obtain the desired community partition. These communities are referred to as functional communities (FCs). This procedure is illustrated in the right panel of Fig 5.

### Synchronization patterns

Once the FCs are identified, we can study how these communities synchronize in response to the stimulation of a specific neuron. To quantify this, we use the Kuramoto order matrix *R* as defined above in Eq. (7). If there are six FCs, this yields a 6 × 6 Kuramoto order matrix, which is always symmetric. An example is shown in Fig 6. Although this provides a useful indication of how strongly pairs of FCs are synchronized, it does not clearly indicate if multiple FCs respond in synchrony or not making it more challenging to identify and quantify patterns of partial synchronization. One way to tackle this is to follow the approach detailed in [49]: First, we apply a threshold *R*_*th*_ to the Kuramoto order matrix, which binarizes the matrix. We then apply the generalized Louvain algorithm for community detection [63, 64] to this binarized matrix. This method assigns an index to each community: if two or more communities share the same index, they form a synchronized group, whereas a community whose index is not shared with any other community is considered asynchronous. The method (which we refer to as the “Louvain synchronous community detection”) is pictorially illustrated in Fig 6. We assign red colors to the synchronized communities and blue colors to the desynchronized ones. Based on the overall synchronization pattern, we classify the system as follows: the pattern is termed *synchronous (sync)* if all communities share the same index (and thus the same color); it is termed *asynchronous (async)* if all communities have distinct indices (or distinct colors). Any intermediate case is classified as a *mixed* or *chimera* pattern, defined as a dynamical state where coherent (synchronized/in-phase) and incoherent (desynchronized/out-of-phase) dynamics coexist simultaneously within the same network of identical or similar units [51]. The pattern shown in Fig 6 is an example of a chimera pattern. The same procedure can be applied to the behavioral circuits by stimulating neurons belonging to different circuits to analyze the resulting synchronization patterns. It is important to note that these patterns depend on the choice of the threshold *R*_*th*_ such that one needs to consider a range of values to establish the robustness of any findings.

**Fig 6.**
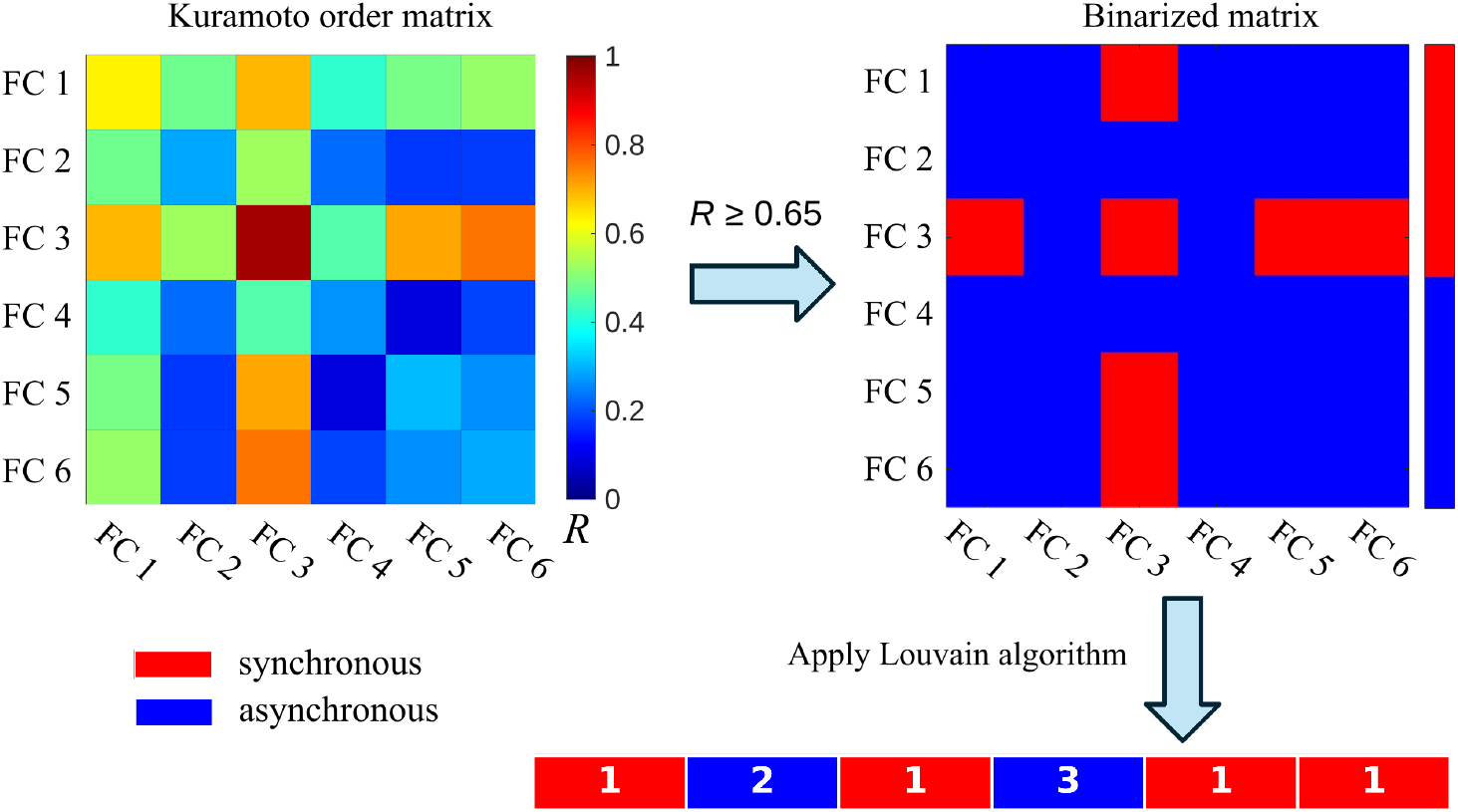
Synchronization pattern detection using the Louvain algorithm. Left: Functional community–based Kuramoto order matrix for a representative case of neuronal stimulation with external current where the stimulated neuron belonged to FC 2. Right: A synchronization threshold of *R*_*th*_ = 0.65 is used to binarize the order parameter matrix, after which the generalized Louvain algorithm is applied to identify synchronized and desynchronized groups. Red denotes synchronized groups, and blue denotes desynchronized ones. The resulting pattern in this case is a chimera pattern.

## Results

Our study aims to understand the relationship between structural connectivity, functional connectivity and biological behavior in *C. elegans* in the context of our model. We begin by investigating how the anatomical network structure shapes global synchronization across neuronal responses. Because these responses are used to identify FCs, this analysis also provides insight into how network structure shapes function.

Next, we focus on the FCs identified by the WSBM method and elucidate the emergent meso-scale functional architecture. Then, we provide a detailed comparison between the FCs and the behavioral circuits. Finally, we analyze the synchronization patterns within both the FCs as well as in the behavioral circuits.

### Effect of structural connectivity on synchronization

We explore the relationship between global synchronization, quantified by the order parameter *Z*, and the underlying structural connectivity of neurons. We collect all neuron stimulation cases with varying current amplitudes and stimulate the *susceptible* neurons. These are defined as neurons that can induce oscillatory responses in other neurons when driven with suitable current amplitudes. The external current and the corresponding susceptible neuron together form a current-neuron pair. We then measure the time-averaged global Kuramoto order parameter *Z*. For each case, we also record the gap junction degree, as well as the in-degree and out-degree of chemical synapses of the stimulated neuron. To establish to which degree the specific structural connectivity is responsible for the emergent global synchronization, we also consider different versions of randomized conectivities. As the gap junction degree of the stimulated neuron has the dominant effect on the global Kuramoto order parameter *Z* as we show below, we consider specifically the following variants of the structural connectome where *only* the gap junction degrees are randomized: (i) a randomly rewired network in which each neuron’s gap junction degree is preserved; (ii) a network with the same gap junction degree distribution, but with node degrees and connections randomly reassigned; and (iii) an Erdős–Rényi–type random network with the same total number of gap edges.

The relationship between the global synchronization *Z* and the gap junction degree of the stimulated neurons, for each of these randomized variants as well as for the original network, is shown in Fig. 7. For the original network, the global synchronization tends to decrease as the gap junction degree of the stimulated neuron increases. The global synchronization order parameter attains its largest mean value when the gap junction degree of the stimulated neuron is zero, i.e., when the stimulated neuron is gap junctionally isolated from the rest of the network. When we consider the different randomized connectivity variants, the largest mean global synchronization always occurs at zero gap junction degree. These observations indicate that stimulation of gap junctionally isolated neurons in the vast majority of cases elicits a global in-phase neuronal response, largely independent of the network architecture. As we show later, these gap junctionally isolated neurons are also captured in our functional community detection and form a separate functional community of their own.

**Fig 7.**
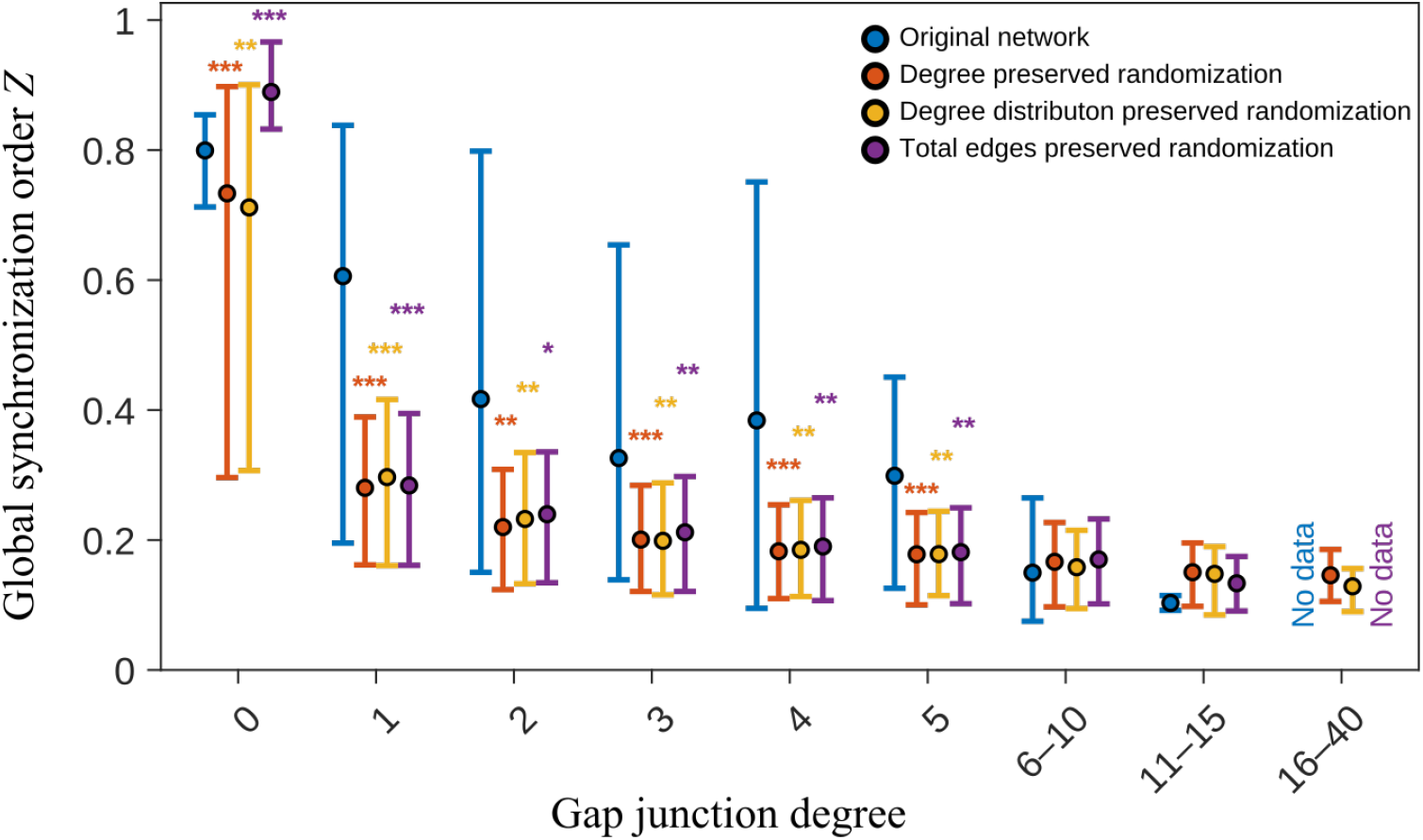
Relation between the global synchronization order *Z* and gap junction degree of the stimulated neuron: The original network and the ensemble averages of its three randomized variants are shown. The dot denotes the mean value and the lower and upper edges represent the 10th and 90th percentiles, respectively. To compare the original network with the random variants, *p* values are calculated by using the two-sided Wilcoxon rank sum test where *, **, and *** denote *p <* 0.05, 0.01, and 0.001, respectively.

Focusing on the nonzero gap junction degrees in Fig. 7, we find that, for degrees less than six, the mean of the global synchronization order parameter is smaller in each of the three randomized variants compared to the original network. Within the same gap junction degree range (1 to 5), the mean value of *Z* decreases with increasing gap junction degree for each randomized variant. This trend also holds for the original network, except for a small increase in the mean value of *Z* at gap junction degree = 4. In all cases with gap junction degree between 1 and 5, however, the distribution of *Z* for each of the three network variants is statistically significantly different from that of the original network, with *p <* 0.05 and lower. It is worth noting that the original network contains only 163 current-neuron pairs (and thus 163 associated gap junction degrees in total), whereas each randomized network variant considers 10 realizations and includes, on average, close to 400 current-neuron pairs per realization. The mean value of *Z*, averaged over all current-neuron pairs, is 0.50 for the original network, 0.22 for the degree-preserved randomization, 0.22 for the degree-distribution-preserved randomization, and 0.21 for the total-edges-preserved randomization. These results indicate that randomizing the connectivity of the gap junctions not only typically increases the number of neurons that are susceptible to external current stimulation (more than double for the randomized realizations compared to the original network), but at the same time the emergent oscillations tend to be more out-of-phase. Another key observation is that, in the original network, the maximum gap junction degree among the susceptible neurons is 14. For the non-Erdős–Rényi–type randomizations, this maximum increases substantially, reaching 40 (the largest gap junction degree present in the original network), see Fig. 7 where data for gap junction degrees greater than 5 are grouped into three categories due to limited sample size and reduced variability. The mean value of *Z* is always very low in this range (when it exists) for all randomization variants and the original network. One possible biological interpretation of these observations is that the native gap junction architecture constrains the susceptibility of neurons, particularly highly connected ones, to externally induced oscillations, thereby limiting excessive reactivity and supporting stable internal control of network dynamics.

### Functional communities (FCs)

Building on our analysis of global synchronization among neuronal responses and its dependence on structural connectivity, we now turn to the functional connectivity between neurons. In our case, when a neuron is stimulated, we focus on the voltage response of the neuronal system and use these responses to compute the pairwise correlations, which we treat as measures of functional connectivity. At the level of individual neurons, these functional relationships reveal whether the responses of two neurons are correlated. However, our primary interest lies in identifying groups of neurons that respond similarly when another neuron is stimulated. This is why we study the meso-scale architecture of the neuronal network by partitioning it into disjoint modules or communities based on functional connectivity. The procedure used to determine these FCs from the neuronal voltage response data is described in the Materials and methods section.

The WSBM approach used for this purpose identifies latent community structures in networks through a generative modeling framework that jointly accounts for edge existence and edge weights, grouping nodes with similar patterns of weighted connectivity across the network. Because the algorithm is non-deterministic, repeated runs typically yield slightly different partitions. To obtain a robust community structure, we perform multiple runs of the WSBM procedure on the co-occurrence matrix, align them, and determine the final assignment of each neuron by taking the mode of its community labels across all runs. For the results presented here, we fix the number of communities to six and perform fifty runs of the WSBM procedure. As a systematic study of varying the number of communities shows (see, e.g., Fig. E in S1 Appendix), this provides the best compromise between capturing sufficiently fine details and ensuring the robustness of the findings. We observe two dominant classes of community assignments emerging from the WSBM approach when the number of communities is set to six. Across multiple runs of the WSBM, type I appears in approximately 70% of the runs, while type II occurs in the remaining 30%. Below, we elaborate on the characteristics of these two community partition types.

- WSBM type I: After arranging the communities by size, we label them as FC 1 (75 neurons), FC 2 (67 neurons), FC 3 (52 neurons), FC 4 (45 neurons), FC 5 (27 neurons), and FC 6 (13 neurons). Examining the gap junction connectivity reveals that these connections are predominantly intra-community rather than inter-community, indicating that the communities are assortative with respect to gap junction. In contrast, chemical synapses are distributed both within and across communities. The gap junction and chemical synapse connectivity patterns among the FCs are shown in Fig 8A.
- WSBM type II: FC 4, FC 5, and FC 6 remain unchanged in WSBM type II when we compare them to type I. The other three communities lose some neurons and gain a few others while preserving their core structure. In this case, FC 1, FC 2, and FC 3 contain 53, 84, and 57 neurons, respectively. We label these communities based on their largest overlap with the corresponding WSBM type I communities. Similar to type I, the FCs in type II are assortative with respect to gap junction connectivity, whereas chemical synapses occur both within and across communities (see Fig 8B).

**Fig 8.**
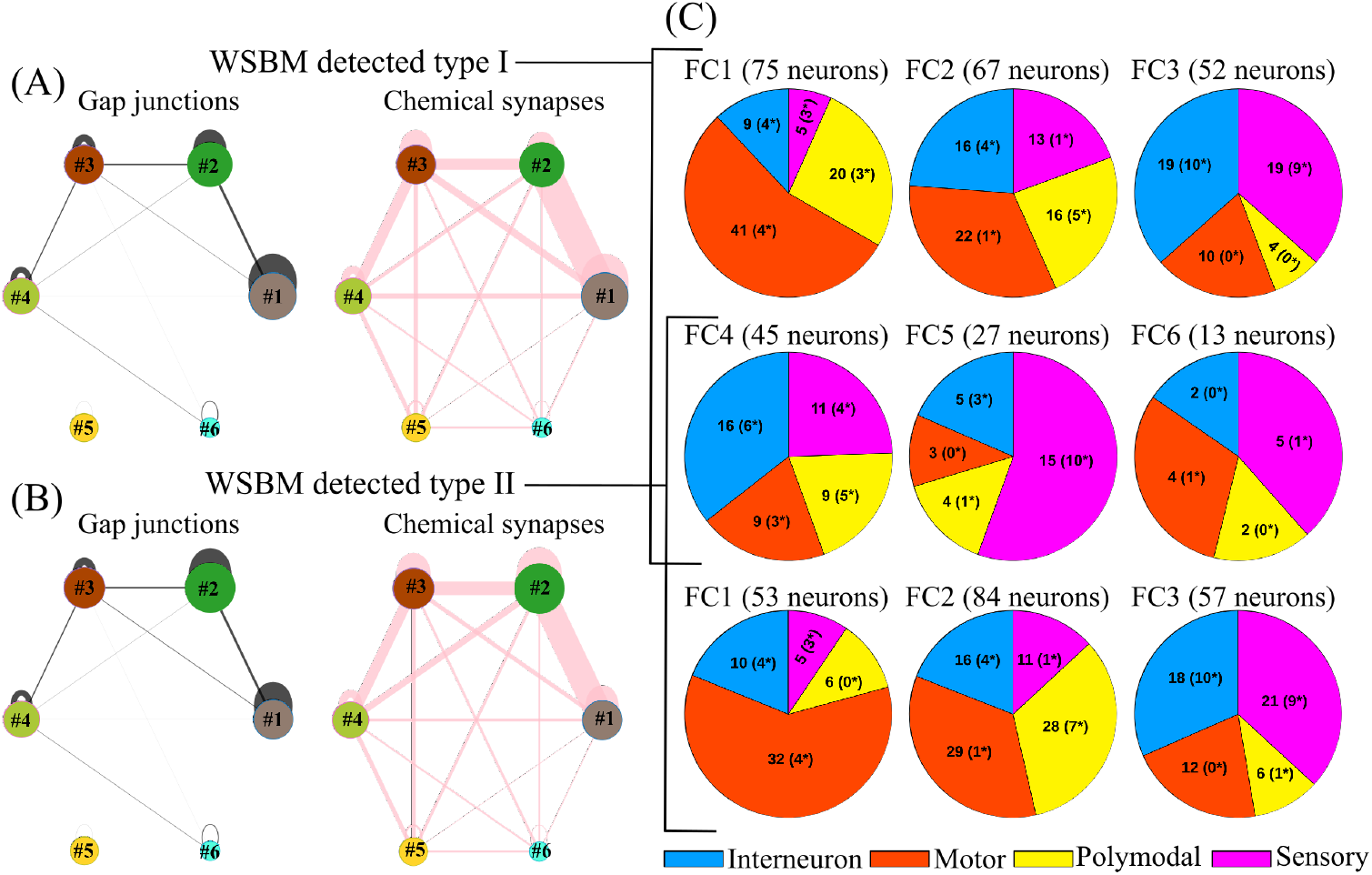
Functional community architecture of *C. elegans* derived from our model. Panels (A) and (B) show the gap junction and chemical synapse connectivity between FCs for WSBM type I and type II, respectively. Gap junctions are predominantly intra-community, whereas chemical synapses are distributed across both intra- and inter-community connections. Panel (C) presents community-wise pie charts illustrating the proportions of sensory, inter-, motor, and polymodal neurons within each community. The numbers in parentheses indicate the count of susceptible neurons in each group. Across all five current amplitudes used, a total of 78 such unique neurons are identified.

We observe that although the neuronal composition of the first three communities differs slightly between type I and type II, the overall pattern of connectivity among the communities remains largely consistent. In both cases, gap junctions are predominantly concentrated within communities rather than across them, whereas chemical synapses are not restricted to within-community connections. Once we have this knowledge, an important question arises: what features of the structural network and the computational model drive this particular form of community partitioning in the first place? The explanation lies in the structure of the gap junction connectivity and how it is encoded in the model. In Eq. (1), neuronal membrane voltages are coupled diffusively through weighted gap junctions, which promotes homogenization of the voltages across connected neurons. Consequently, stronger gap junction coupling leads to stronger correlations in their membrane voltages. When these voltage correlations are used as functional connectivity measures for community detection, neurons with stronger mutual gap junction connections naturally cluster together. In contrast, chemical synapses do not promote synchronization of neuronal voltages, and therefore, their connections are distributed both within and across communities. This mechanism accounts for the grouping patterns and their connectivity observed in both types of community partitions detected by the WSBM procedure.

It is important to note that FC 5, which appears identically in both types of partitioning, is gap junctionally isolated from the rest of the communities. Out of the 27 neurons in this community, 25 neurons have no gap junction connections with any others, and there is a pair of neurons that are mutually connected. To put this in perspective relative to the full gap junction connectivity, the structural network consists of a giant connected component of 248 neurons, two smaller components of 2 and 3 neurons, and 26 isolated neurons. Thus, FC 5 contains nearly all of the neurons that are gap junctionally isolated. This community also contains the largest proportion of sensory neurons (16 out of 27), and, as we will show below, stimulation of neurons within this community produces the most strongly synchronized, community-wide response pattern. This is consistent with our earlier analysis of the relationship between structural connectivity and global synchronization, where we found that a zero gap junction degree of the stimulated neuron corresponds to the highest global synchronization.

While the different neuronal types are not captured in our model equations (Eqs. (1)-(2)), their role might be partially encoded in the underlying structural connectome and, hence, in the arising FC. The community-wise composition of different neuronal types in both types of partitions is visualized through the pie charts shown in Fig 8C. Once again, we observe strong similarity between the two. In both cases, FC 1 is composed primarily of motor neurons. A substantial number of interneurons are present in FC 1, FC 2, FC 3, and FC 4. Most sensory neurons are grouped in FC 3, FC 5, and FC 2. FC 6 has the smallest community size and contains only 13 neurons in total. The rich-club neurons are primarily located in FC 1, FC 2, and FC 3 for both types of partitioning. The distributions of rich-club neurons [65] across the FCs for both partition types are presented in Table B of S1 Appendix.

### Comparison of FCs with behavioral circuits

One of the central objectives of this study is to understand how the behavioral circuits in *C. elegans* relate to functional connectivity, which in turn we have shown to be largely shaped by the underlying gap junction structural connectivity in the context of our model. The neuronal circuits associated with five specific *C. elegans* behaviors are described in the Materials and Methods section. Locomotion, characterized by undulatory body movements, is classified into forward and backward modes depending on the direction of motion. The other behaviors correspond to different forms of sensory processing: chemosensation (odor, taste etc.), mechanosensation (gentle or harsh touch), and klinotaxis (navigation in salt gradients).

To reveal how these behaviors are functionally interconnected, Table 3 presents a comparison between the FCs and the behavioral circuits, showing how the neurons in the behavioral circuits map onto the different FCs. The FCs are mutually exclusive and exhaustive, comprising all 279 neurons, whereas the behavioral circuits are not exclusive and together include a total of 123 neurons. The remaining 156 neurons are not part of any of the behavioral circuits studied in this paper. Earlier, we demonstrated a strong similarity between the FCs in type I and type II. However, their differences become evident when we examine how neurons from the behavioral circuits are distributed across the FCs. In type I, most neurons involved in forward locomotion (22 out of 24) and backward locomotion (23 out of 31) are grouped into FC 1. In contrast, in type II, the majority of forward locomotion neurons shift to FC 2 (now 20 out of 24 neurons are in FC 2), while the backward locomotion neurons remain in FC 1. The distributions for the other behavioral circuits remain largely consistent across both types. As noted earlier, FC 4, FC 5, and FC 6 are identical in both partitions, so no differences are observed there. Thus, the key distinguishing feature between the two types of community partitions is whether the locomotor neurons are grouped within a single community (type I) or separated across different communities (type II) — the latter case becoming more prevalent if the number of functional communities is increased (see Fig. E of S1 Appendix).

**Table 3.** Functional community-wise distribution of the behavioral circuits.

| Functional Communities | Type | Behavioral Circuits |  |  |  |  |
| --- | --- | --- | --- | --- | --- | --- |
|  |  | Forward Locomotion | Backward Locomotion | Chemo-sensation | Mechano-sensation | Klino-taxis |
| **FC 1** | I (II) | 22 (4) | 23 (23) | 6 (6) | 9 (7) | 0 (1) |
| **FC 2** | I (II) | 2 (20) | 3 (3) | 5 (6) | 4 (6) | 8 (8) |
| **FC 3** | I (II) | 0 (0) | 5 (5) | 14 (13) | 14 (14) | 4 (3) |
| **FC 4** | I & II | 0 | 0 | 9 | 8 | 8 |
| **FC 5** | I & II | 0 | 0 | 6 | 2 | 2 |
| **FC 6** | I & II | 0 | 0 | 6 | 1 | 1 |

The table shows how neurons from each behavioral circuit are distributed across the identified functional communities. Values outside parentheses correspond to Type I, while values inside parentheses correspond to Type II communities, as detected using WSBM.

Another important point to understand is how distributed or concentrated the behavioral circuits are across the FCs. The forward and backward locomotion circuits are largely confined to a single FC, indicating a high degree of concentration. In contrast, the chemosensation circuit is spread across all six FCs (in both types), with its largest core located in FC 3. A similar pattern is observed for the mechanosensation circuit, although it shows only a small overlap with FC 6. The klinotaxis circuit is also distributed, spanning five FCs in type I and all six FCs in type II. Thus, compared to the locomotor circuits, which are highly localized, the sensory circuits (chemosensation, mechanosensation, and klinotaxis) are more widely distributed. Looking from the opposite perspective, the first three FCs contain neurons from all behavioral circuits (with FC 3 containing neurons from backward locomotion only). In contrast, the last three FCs include neurons exclusively from the sensory-driven circuits – chemosensation, mechanosensation, and klinotaxis.

### Synchronization patterns in FCs

When a neuron is stimulated, it generates a system-wide response mediated through gap junctions and chemical synapses. Having already identified the FCs based on voltage-response correlations, we now examine how the FCs synchronize among themselves when the neurons of each FC are stimulated. For this purpose, we compute the inter-community Kuramoto order matrix *R* for each neuron stimulation case in every FC including all five current amplitudes (see Materials and Methods for details) and then calculate the average Kuramoto order matrix. Here, we present results only for the FCs in type I. Corresponding results for type II, which are qualitatively very similar, are provided in the Supplementary Materials (Fig. C, D in S1 Appendix).

There are 14 susceptible neurons in FC 1 that elicit oscillatory responses when stimulated with specific external currents. In total, 19 current–neuron pairs in FC 1 generate such oscillatory behavior. For the remaining FCs, these numbers are: FC 2: 11 neurons and 17 current–neuron pairs, FC 3: 19 neurons and 37 pairs, FC 4: 18 neurons and 38 pairs, FC 5: 14 neurons and 49 pairs, FC 6: 2 neurons and 3 pairs. For each current–neuron pair, we compute a Kuramoto order matrix. We then average these matrices across all pairs within a given FC to obtain a single representative Kuramoto order matrix for that community. In this way, each FC is associated with one averaged Kuramoto order matrix reflecting the “typical” synchronization pattern elicited by its susceptible neurons.

In Fig 9, we show the averaged Kuramoto order matrices for the six FCs in type I. When neurons in FC 1, FC 2, and FC 3 are stimulated, the overall level of synchronization between FCs is relatively low (see panels A-C of Fig 9), whereas stimulation of neurons in FC 4, FC 5, and FC 6 results in higher synchronization (see panels D-F of Fig 9). The highest level of synchronization is observed when neurons in FC 5 are stimulated (Fig 9E). However, in all cases, FC 5 exhibits the lowest level of synchronization with the other FCs (fifth row and column in each panel of Fig 9). In other words, FC 5 elicits the most synchronized response overall, yet fails to synchronize effectively with the other communities.

**Fig 9.**
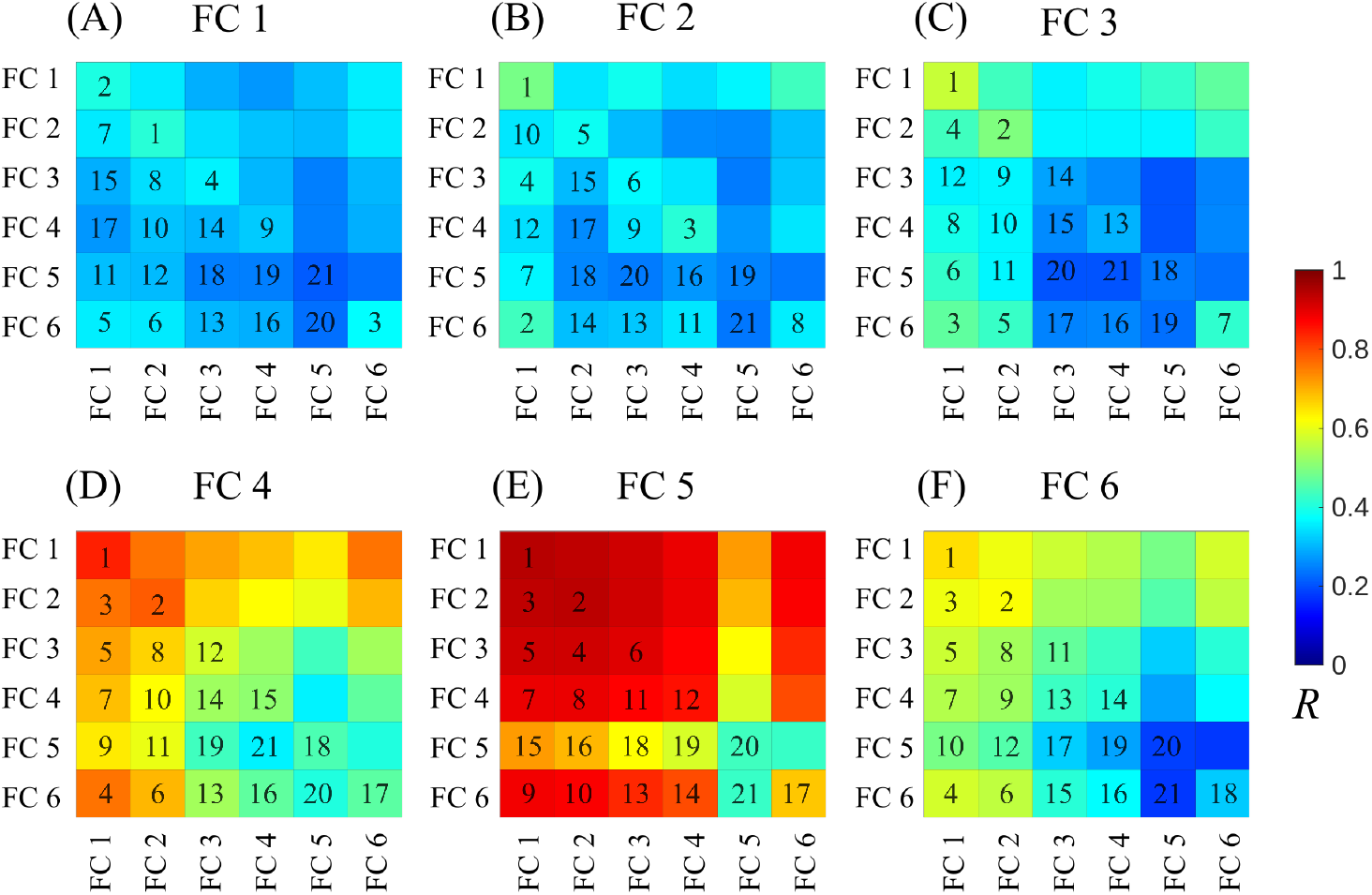
Averaged Kuramoto order matrix *R* for stimulation applied to neurons in different FCs (type I). Panels (A)-(F) correspond to stimulation of neurons in FC 1 through FC 6, respectively. The colorbar indicates the value of *R*, reflecting the degree of synchronization between pairs of FCs. The rank numbers in each matrix are assigned in decreasing order of *R*, where 1 denotes the highest level of synchronization and 21 denotes the lowest. Stimulation of neurons in the first three FCs results in relatively low synchronization across communities, whereas stimulation of neurons in the last three FCs produces higher overall synchrony. In all cases, FC 5 consistently shows the lowest level of synchronization with the other communities including itself.

This is expected based on our findings above. As noted previously, FC 5 is gap junctionally isolated from the rest of the FCs and connects to them only via chemical synapses. It also has very few gap junction connections among its own neurons. In fact, of the 14 susceptible neurons in this community, 12 have no gap junctions with any other neurons, and the remaining two are connected only to each other. Since gap junction connections promote synchronization of neuronal voltages through diffusive coupling, it is expected that neurons in FC 5, having the fewest gap junctions, exhibit the lowest level of synchronization. Similarly, the reason why other communities become strongly synchronized when FC 5 neurons are stimulated lies again in the gap junction connectivity. Indeed, as we have shown before, neurons with a zero gap junction degree tend to elicit the highest level of synchronization in the network.

The fact that FC 5 induces the strongest synchronized response, and that FC 1 and FC 2 exhibit the highest levels of synchrony, is also reflected at an individual neuron stimulation level when we inspect the synchronization patterns through the lens of Louvain synchronous community detection method (as described in the Materials and Methods section). When a neuron is stimulated, this method determines whether two FCs are synchronous based on a predefined synchronization threshold *R*_*th*_ (see Fig. B in S1 Appendix where we vary *R*_*th*_). We perform this analysis for all current–neuron pairs in all FCs, and the resulting patterns are summarized in Fig 10. In panel (a), these patterns are grouped according to the FC being stimulated. When stimulated, FC 5 produces a fully synchronized response (100% sync patterns), whereas FC 1 yields the most desynchronized response (80% async patterns), followed by FC 2 (69.2% async patterns). FC 3, in contrast, exhibits the highest proportion of mixed responses, with approximately 54% of its outcomes classified as chimera patterns (combining all mixed types). At the same time, the large number of different chimera patterns further indicates that FC-3 is endowed with a large dynamical flexibility, meaning the various stimuli can lead to different responses. FC 4 is the only community whose stimulation produces all three types of synchronization patterns (each greater than 10%), with approximately 41% sync, 23% async, and 36% chimera responses. FC 6 consistently produces a fixed chimera response for each stimulation; however, it contains only 13 neurons and 3 current–neuron pairs, making it of limited interest.

**Fig 10.**
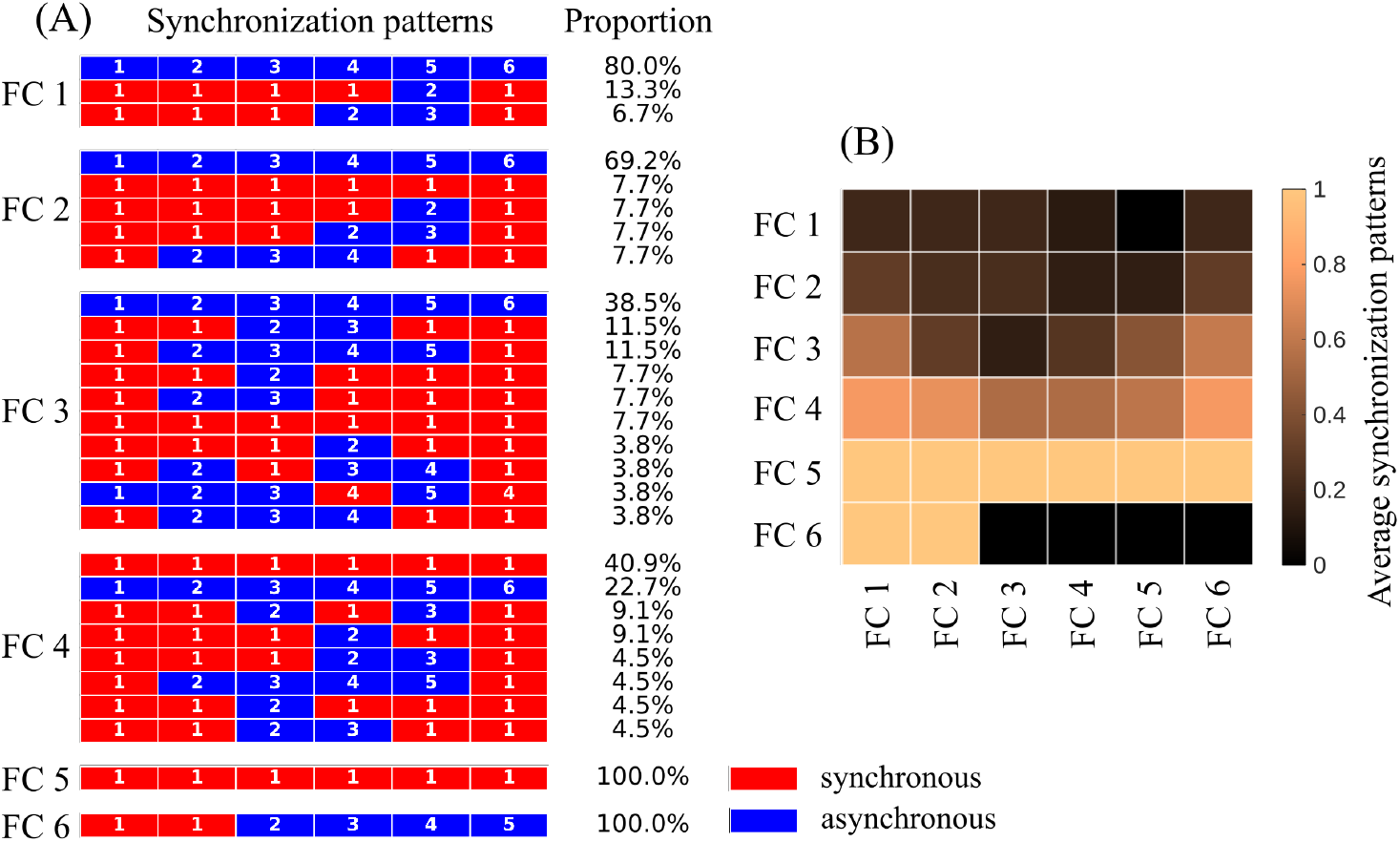
Emergent synchronization patterns following FC-wise (type I) stimulation. (A) Synchronization pattern types and their proportions for each FC when its neurons are stimulated. Each horizontal panel inside each FC represents a synchronization pattern; see Fig. 6 and the accompanying text for details. (B) Average synchronization patterns: in the 6×6 matrix, the *i,j*-th entry indicates the probability that the *j*-th FC becomes part of a synchronous group when the *i*-th FC is stimulated where *i* and *j* indicate the rows and columns, respectively. Here, we use *R*_*th*_ = 0.6.

The information provided in Fig 10A can be represented more compactly by measuring the probability that a given FC belongs to a synchronous group (indicated in red) when neurons of another FC are stimulated. These probabilities are computed and presented in Fig 10B, where the *i,j*-th entry of the matrix represents the probability of the *j*-th FC of being a member of a synchronized group when the neurons of the *i*-th FC are stimulated. In other words, the matrix captures the column-wise response to row-wise stimulation. It also recapitulates the key observations: FC 5 drives the strongest synchronized response, while FC 1 and FC 2 exhibit the highest levels of synchrony when stimulated.

### Synchronization patterns in behavioral circuits

Having established the properties of the emerging synchronization patterns in the FCs, we now turn to the synchronization patterns when viewed from the perspective of the behavioral circuits. In the forward locomotion circuit, there are 24 neurons, of which only 2 are susceptible neurons that generate oscillatory responses upon stimulation.

Each of these neurons produces oscillations for just one of the five tested current amplitudes, resulting in only 2 current–neuron pairs for this circuit. For the other behavioral circuits, the numbers are as follows: backward locomotion contains 6 susceptible neurons with 8 current–neuron pairs; chemosensation has 18 susceptible neurons and 39 pairs; mechanosensation includes 14 susceptible neurons with 29 pairs; and klinotaxis comprises 5 susceptible neurons with 13 current–neuron pairs.

We stimulate the neurons in each behavioral circuit and compute the Kuramoto order matrix *R* between the circuits (see Materials and Methods for details). These values are averaged over all current–neuron pairs within each circuit, resulting in five averaged Kuramoto order matrices corresponding to the five behavioral circuits. The matrices are shown in Fig 11. Apparently, stimulation to the forward and backward locomotion circuits produces a comparatively lower level of synchronization than stimulation to the chemosensation, mechanosensation, and klinotaxis circuits (by stimulation of a given circuit, we specifically mean stimulation of the susceptible neurons within that circuit). Another notable observation is that the forward and backward locomotion circuits exhibit the highest level of synchronization across all circuit stimulations (first two rows and columns in each panel of Fig 11). In contrast, the chemosensation circuit shows the lowest level of synchronous response (third row and column in each panel of Fig 11). These observations are not as pronounced in Fig 11A, as it includes only two neuron stimulation cases as mentioned above.

**Fig 11.**
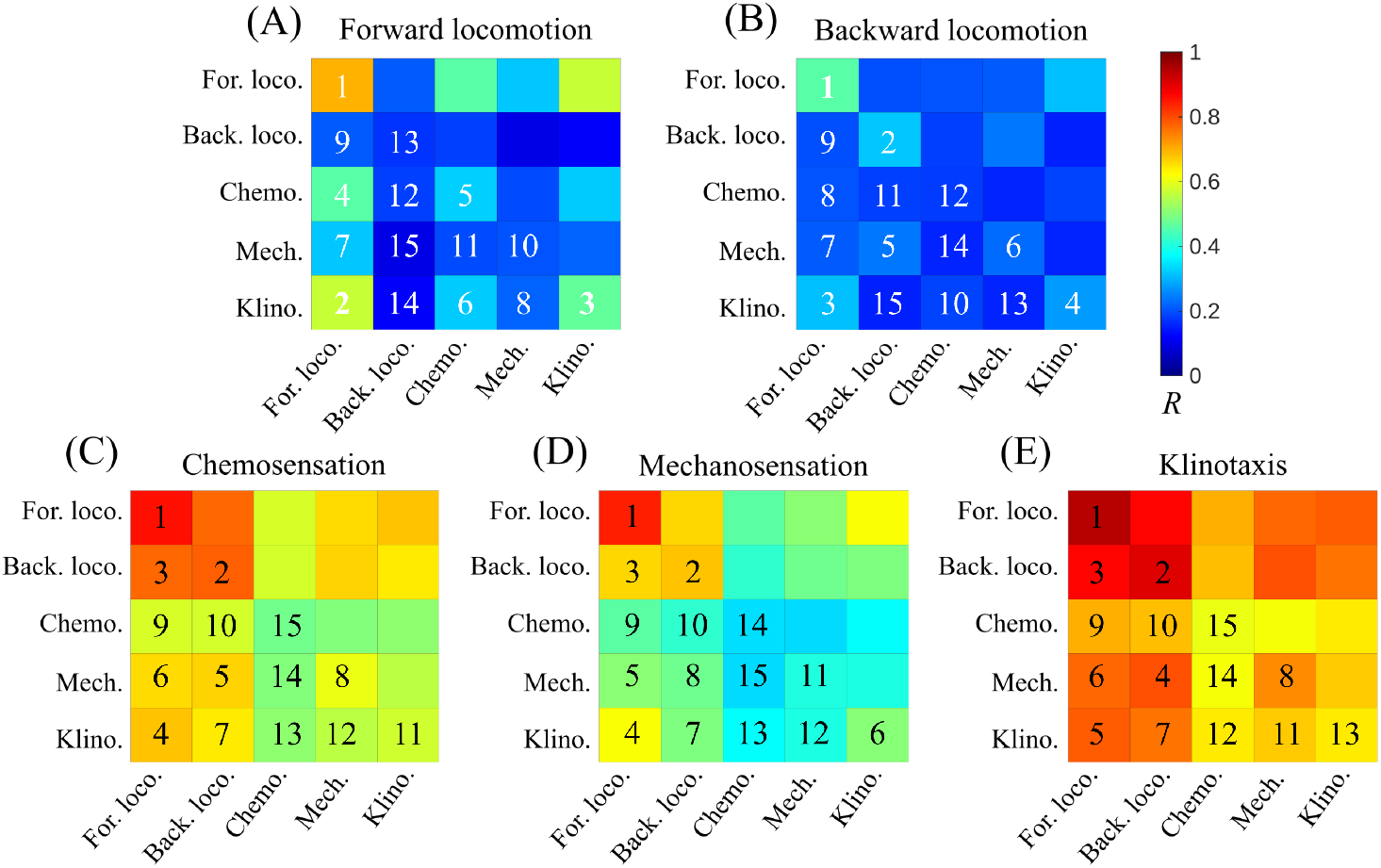
Averaged Kuramoto order matrix *R* for stimulation of neurons in the behavioral circuits. Panels (A)–(E) correspond to stimulation of neurons in the forward locomotion, backward locomotion, chemosensation, mechanosensation, and klinotaxis circuits, respectively. The rank numbers in each matrix are assigned in decreasing order of *R*, where 1 denotes the highest level of synchronization and 15 denotes the lowest. Stimulation to the neurons of the locomotion circuits produces a low level of synchronization, whereas comparatively higher synchronization is observed when neurons in the chemosensation, mechanosensation, and klinotaxis circuits are stimulated.

We can now revisit the observations of the emergent synchronization patterns from the FC perspective and relate them to the behavioral circuits. Stimulation of FC 1 and FC 2 produced low levels of synchronization, similar to the responses observed for forward and backward locomotion stimulation. From Table 3, we see that FC 1 and FC 2 contain the largest number of neurons from the two locomotion circuits.

Conversely, forward and backward locomotion circuits exhibit the highest levels of synchronous responses, mirroring the behavior of FC 1 and FC 2. In contrast, establishing a direct correspondence for the chemosensation, mechanosensation, and klinotaxis circuits is less straightforward, as these circuits are more distributed across multiple FCs compared to the concentrated locomotion circuits. From the synchronization patterns in Fig 11, we can infer that stimulating a behavioral circuit distributed across several FCs tends to produce a more synchronized response than stimulating a circuit concentrated in just a few FCs.

Next, we scrutinize the synchronization patterns in the behavioral circuits at the level of individual neuron stimulation. Using the Louvain synchronous community detection procedure described in the Materials and Methods, we determine whether a stimulated neuron causes a circuit to be synchronously grouped with other circuits. By aggregating all circuit-wise neuron stimulation cases, the results are presented in Fig. 12. There is a clear predominance of desynchronized patterns for stimulation within the locomotion circuits (in forward locomotion, 50% async, 50% chimera). In contrast, all three synchronization patterns – sync, async, and chimera – are observed for stimulations in the chemosensation, mechanosensation, and klinotaxis circuits, with the highest proportion of synchronous patterns occurring during klinotaxis stimulation.

**Fig 12.**
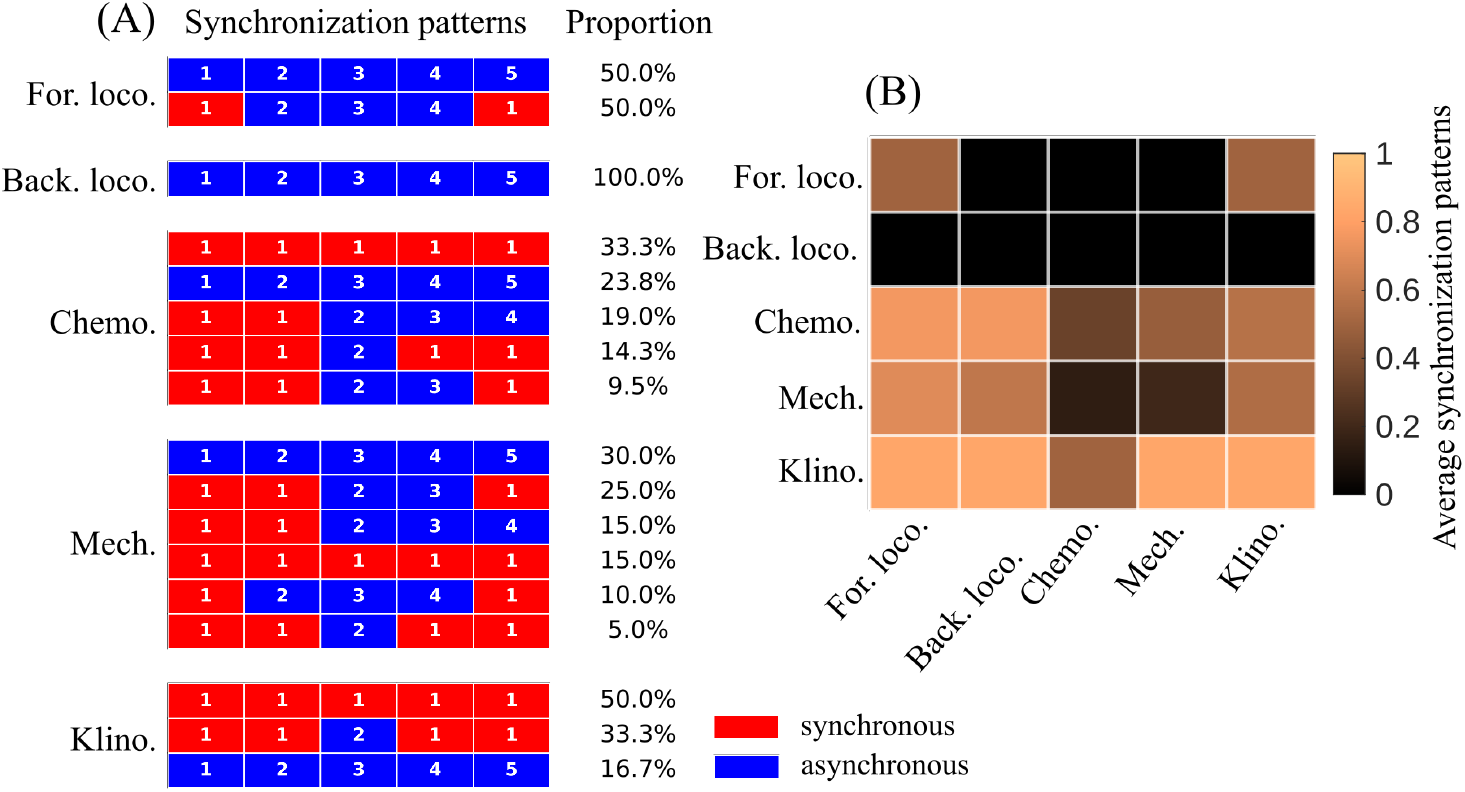
Emergent synchronization patterns for stimulation of the behavioral circuits. (A) Patterns of synchronization and their proportions in each behavioral circuits when its neurons are stimulated. Each horizontal panel inside each behavioral circuit represents a synchronization pattern; see Fig. 6 and the accompanying text for details. (B) Corresponding average synchronization patterns. It denotes the probability of a circuit being a member of a synchronous group when some other circuit is stimulated. Here, we use a synchronization threshold *R*_*th*_ = 0.6.

Across all behavioral circuit stimulation conditions, the chemosensation circuit exhibits the lowest level of synchrony. From the synchronization patterns in Fig. 12A, we observe that the chemosensation circuit is either synchronized with all other circuits or remains desynchronized; the latter case dominates, indicating that the chemosensation circuit is often not part of the synchronized group. This is also reflected in the average Kuramoto order matrices in Fig. 11, where the weakest synchronization appears along the third row and column corresponding to the chemosensation circuit. Stimulation of the mechanosensation circuit yields the largest number of synchronization patterns and the highest fraction of chimera patterns (55%) with the largest diversity of chimera types (four) indicating a high degree of dynamical flexibility. Upon stimulation of other circuits, it also becomes synchronized with a varying probability (see the fourth column of Fig. 12B).

## Discussion

Anatomical connectivity (the connectome) provides the structural constraints [52] – the “wiring diagram” that bounds which pathways can, in principle, transmit and transform signals – whereas functional connectivity captures state-dependent interactions that actually shape neural dynamics during behavior [23, 66] (e.g., strengths, signs, and temporal properties that are not determined by anatomy alone). In practice, behavior depends on both [25]: the same anatomical scaffold can support different dynamical regimes and stimulus responses, so comparing anatomical and functional connectivity is essential for identifying which structural motifs are actually recruited, and how. For the same reason, models and experiments must be pursued in parallel: experiments provide whole-brain measurements and targeted perturbations to define and validate effective interactions [57], while models convert those constraints into mechanistic hypotheses, sharpen causal predictions, and expose which additional measurements are necessary to disambiguate mechanisms [25]. Using an experimentally motivated computational model, we investigated how structural connectivity shapes functional properties and how these, in turn, relate to specific *C. elegans* behaviors.

### Linking structure, function, and behavior in *C. elegans*

Although previous experimental studies have reported a weak relationship between anatomical and measured functional as well as signaling connectivity [11, 12], other work has reached seemingly contrasting conclusions, reporting causal interactions between the two [13, 17]. Nevertheless, it is broadly accepted that behavior is both a structural and functional property [16, 23, 25]. It emerges from the functional dynamics of neural circuits operating under the constraints imposed by the underlying structural architecture. We considered several well-known behavioral circuits and compared them with our FCs by examining their neuronal composition and synchronization patterns. The FCs are inferred from correlations between model-derived neuronal voltage responses, which are shaped by the underlying structural connectivity among neurons.

### Structural influence on FCs

Gap junctionally isolated neurons always emerged to be a key factor for a high level of global synchronization and eventually had an important role in the FC formation. A clear trend has been observed in which the global synchronization order *Z* is high when the gap junction degree of the stimulated neuron is zero or very low, and decreases as the degree of the gap junction increases (see Fig. 7). It is also true that neurons with no gap junctions have the highest probability of eliciting global oscillations in the first place (i.e., of being susceptible), whereas neurons with very high gap junction degree are rarely susceptible (see Fig. J of S1 Appendix). In contrast, we do not observe any clear trend for chemical synapse in-degree or out-degree, either in susceptibility (Fig. J of S1 Appendix) or in the propensity to elicit globally in-phase responses (see Fig. H in S1 Appendix). This disparity is also evident in the gap junction and chemical synapse connectivity among the FCs in Fig. 8: Gap junction connections are largely confined within FCs (intra-community), whereas chemical synapses extend both within and across FCs (intra- and inter-community). In earlier studies, gap junctions have been reported to play a crucial role in *C. elegans* behaviors like locomotion [67], and in some cases dominate certain levels of sensory processing like the ‘hub-and-spoke’ pattern for the convergence of multiple head sensors to communicate via gap junctions onto a single interneuron [68]. On the other hand, sensory representations during olfactory responses have been shown to be independent of the chemical synaptic connections [69]. In our study, gap junctions emerge as a crucial factor for collective synchronous response, consistent with previous findings that demonstrated the role of gap junctions in synchronous neuronal activity [70], while the role of chemical synapses, although important for signal propagation, remains less clear.

In the adult hermaphrodite *C. elegans*, neuronal cell bodies follow a stereotyped layout along the body axis, with dense ganglia at the head and tail connected by longitudinal nerve cords [1]. Sensory neurons are mostly positioned at the periphery in the head and tail, where they contact the environment; interneurons are concentrated in central ganglia around the nerve ring and along smaller midbody/tail ganglia, integrating multimodal inputs; and motor neurons are organized in repeating units along the ventral (and dorsal) nerve cords adjacent to body-wall muscles [52]. Together, this arrangement creates a compact but highly ordered sensorimotor architecture linking environmental inputs to motor outputs [71]. When we compare the FCs with the body positions of the neurons, we find that neurons in FC 1 are distributed along the entire body (see Fig. G in S1 Appendix). This spatial spread is somewhat reduced, but still present, in FC 2 (neurons in FC 2 are more prominently distributed over the body in type II than in type I). By contrast, neurons in FC 3, FC 4, FC 5 and FC 6 are predominantly located near the head, with fewer neurons near the tail or midbody. It is well established that locomotor decision-making and control are primarily centered in the head (around the nerve ring), whereas the execution circuitry, notably the motor neurons, is distributed along the ventral nerve cord (and, via commissures, the dorsal nerve cord) down the body [3]. In our study, we found a strong one-to-one correspondence between FC 1 and FC 2 and the locomotion circuits, and this correspondence is further supported by the spatial distribution of the neurons.

### FCs and behavioral circuits

One motivation for focusing on meso-scale functional community structure is to enable direct comparisons between these communities and established behavioral circuits. We find a strong one-to-one correspondence between the forward and backward locomotion circuits and two FCs. In contrast, the chemosensation, mechanosensation, and klinotaxis circuits were distributed over the FCs. These observations are robust with the choice of number of communities. For each choice of number of FCs, comparison with the behavioral circuits showed that the locomotion circuits remained relatively segregated across FCs, whereas the sensory circuits were consistently more distributed. The main effect of changing the number of communities was on the relative frequency of two types of community partitions: type I, in which both the forward and backward locomotion circuits overlap largely with the same FC, and type II, in which the forward and backward locomotion circuits have their maximum overlap with two different FCs. When the number of communities is small, Type I is observed more frequently than Type II. As the number of communities is increased beyond six, Type II becomes more prevalent, which is expected since a larger number of communities can capture even slightly dissimilar dynamical features (see Fig. E in S1 Appendix).

The sensory circuits – chemosensation, mechanosensation, and klinotaxis – are the most suitable targets for stimulation, in the sense that the resulting synchronization patterns are largely synchronous (see the average synchronization patterns in Fig. 12B). On the other hand, the forward and backward locomotion circuits are less favorable stimulation targets for generating in-phase responses or synchronous response patterns (see panel A-B of Fig. 11 and first two rows of Fig. 12B), yet they exhibit the highest levels of synchronization when another circuit is stimulated (see the first two rows and columns of panel C-E in Fig. 11 and first two columns in Fig. 12B). The emergent synchronization patterns include all three types (async, chimera, sync) under stimulation of the sensory circuits (see Fig. 12A), and the same holds for some of the FCs (see FC 2, FC 3 and FC 4 stimulation cases in Fig. 10A where all three synchronization patterns are observed). This demonstrates the dynamical flexibility of the sensory circuits as well as some of the FCs derived from our study. In particular, our observations are consistent with previous studies. For example, the mechanosensation circuit includes neurons that encode gentle touch via touch receptor neurons (TRNs) and harsh touch via nociceptive neurons such as PVD, and it exhibits distinct response patterns depending on the stimulus type [72]. We have also found a large repertoire of response patterns upon stimulation of the mechanosensation circuit (see the fourth panel in Fig. 12A). As another example, it has been experimentally established that the chemosensation circuit comprises diverse sensory neurons tuned to different odorants and tastants and therefore exhibits heterogeneous response patterns [38, 69], whereas motor actions are often governed by comparatively simple, low-dimensional dynamics [10]. This contrast is also reflected in our model simulations: the chemosensation circuit typically exhibits an out-of-phase response with other circuits (see the third column of Fig. 12B) and multiple response patterns upon its stimulation (see the chemosensation stimulation case in Fig. 12A), whereas the responses of the locomotion circuits are mostly in-phase (first and second columns of Fig. 12B) and the number of synchronization patterns is also limited (see the locomotion circuits stimulation cases in Fig. 12A). Anatomically, the sensory circuits are connected to the locomotion circuits along feed-forward pathways [71]. This organization may explain why stimulation of the sensory circuits consistently leads to in-phase responses of the locomotor circuits, whereas the reverse is not observed.

### Comparison with signaling communities

How structural connectivity supports functional repertoire is a central question in biology. In this context, a brain-wide causal “signal propagation atlas” was recently obtained via direct optogenetic activation combined with simultaneous whole-brain calcium imaging [73]. This signaling network, consisting of 188 head neurons (including 20 pharyngeal neurons, which we exclude from our structural connectome data), has also been analyzed for its modular structure by optimally partitioning it into six signaling communities (SCs) [11]. Although a direct comparison is not feasible due to mismatches in neuron number and neuron types, we nevertheless find that the SCs are meaningfully different from the FCs analyzed in this study. Comparisons of these SCs with the FCs and with the behavioral circuits are presented in Table C and Table D of S1 Appendix, respectively. We also investigate the synchronization patterns within the SCs and present the results in Fig. F of S1 Appendix. Factors such as polysynaptic chains and extrasynaptic signaling [74], which contribute to effective connections between neurons, are captured experimentally in the signal propagation atlas and are therefore reflected in the SCs, but they are absent in our model simulations.

Nonetheless, our analysis remains valuable because it links specific *C. elegans* behaviors to FCs derived from a computational model, thereby providing insights in regimes where experimental evidence is still limited. Specifically, although simultaneous population recordings in *C. elegans* are now possible, obtaining comprehensive, brain-wide response patterns across diverse natural behaviors remains experimentally challenging [57, 75].

Our method suggests that if a behavioral circuit spans across multiple FCs, then the whole-brain response upon stimulation to that circuit will be more in-phase compared to the stimulation to a behavioral circuit which is largely segregated to a single FC.

### Linking synchronization patterns to neuron types

In our model, neuron types are not explicitly encoded in the dynamical equations and enter only through the structural connectome. Nevertheless, several features, such as how the identity of the stimulated neuron relates to the resulting response patterns, and which neurons exhibit the strongest in-phase responses, show a strong correspondence with particular neuron types. This motivates a closer inspection of the synchronization features and their possible relationship to neuron type. The sensory circuits contain a substantial fraction of sensory neurons: they constitute 57%, 53%, and 22% of the chemosensation, mechanosensation, and klinotaxis circuits, respectively. The higher synchronization observed upon stimulation of these circuits may therefore be related to their large sensory-neuron composition. Consistent with this, our functional community analysis showed that FC 5, which elicits the highest level of synchronization, also comprises 56% sensory neurons. To examine this more directly, we quantified the global synchronization order parameter *Z* under stimulation of different neuron types (see Fig. I in S1 Appendix). We find that the median value of *Z* is highest for sensory-neuron stimulation (0.71), compared with 0.31, 0.51, and 0.20 for interneurons, motor neurons, and polymodal neurons, respectively. Moreover, an analysis of response patterns (sync, async, and chimera) by using the Louvain synchronous community detection method shows that sensory-neuron stimulation yields the largest fraction of fully synchronized outcomes (see Fig. I in S1 Appendix). Together, these results provide strong evidence that stimulating sensory neurons promotes synchronized neuronal responses. However, it would be incorrect to infer a one-to-one correspondence between sensory-neuron content and a high level of synchronization. For example, FC 3 contains 36% sensory neurons, yet stimulation of FC 3 produces predominantly asynchronous responses (see Fig. 9C and Fig. 10). Conversely, the klinotaxis circuit, despite containing only 22% sensory neurons, exhibits very high synchronization (see Fig. 11E and Fig. 12). One can also observe that in all six averaged Kuramoto order matrices shown in Fig 9, overall FC 1 exhibits the highest level of synchrony, followed by FC 2. These two communities contain the largest proportions of motor neurons – 55% in FC 1 and 33% in FC 2. Similarly, the forward and backward locomotion circuits are made up of 29% and 68% motor neurons, respectively, and they respond most synchronously (see Figs. 11 and 12). Combined with the observation that FC 5 as well as the chemosensation and mechanosensation circuits which are largely made up of sensory neurons, elicit strong synchronization, this again supports the notion of information processing along sensorimotor pathways, which are well established [71, 76]. There may be possible relations between neuron type and structural features such as degree distributions etc. which we showed to affect the synchronization behavior (e.g., zero gap junction leading to high synchronization). Future work could explicitly incorporate neuron-type-specific properties in the model to directly examine how particular neuron types influence synchronization.

### Comparing model simulations and experimental observations

A key limitation of our study is that the mapping between “behavioral circuits” and the underlying biophysical substrates is only approximate, because circuit descriptions in *C. elegans* are inferred from a heterogeneous literature and remain incomplete wherever electrophysiological ground truth is sparse (e.g., uncertainties about synaptic sign and context dependence). In addition, our *in silico* paradigm assumes that the network response to stimulation can be characterized under conditions analogous to whole-brain readout; however, simultaneously measuring all relevant neurons *in vivo* remains technically challenging, and typical whole-brain imaging often trades temporal resolution and signal fidelity for coverage. Relatedly, our simulations use constant external input currents to elicit and sustain oscillatory regimes, but implementing truly constant, cell-specific current injection at scale is generally not feasible experimentally, and common perturbation modalities (optogenetics, mechanostimulation) impose different input statistics and nonlinearities. Finally, while our analysis emphasizes phase relationships (after normalization) and treats amplitude heterogeneity as secondary, experimental recordings may not reliably capture voltage amplitude variations across the entire network, since calcium signals are indirect, neuron-specific, and can saturate or compress dynamics – potentially obscuring amplitude-dependent effects that our model allows in principle.

## Conclusion

Understanding how structural connectivity shapes brain function and behavior remains a fundamental and open question in neuroscience. In this study, we use a computational model to investigate how the structural connectome of *C. elegans* influences functional connectivity *in silico*. Building on this, we further explore how some well-known behaviors and their associated neuronal circuits relate to these underlying structural and functional properties. Our analysis revealed several key characteristics of the behavioral circuits and their relationship to structure and function, motivating the following hypotheses: (i) the locomotion circuits function largely independently yet synchronize robustly: their collective dynamics appear similar regardless of how input signals reach them or how they are stimulated; (ii) sensory circuits comprise multiple functional components and exhibit stronger, stimulus-dependent sensitivity; and (iii) neurons that are gap junctionally isolated are optimal stimulation targets for eliciting a collective synchronous response. Testing these hypotheses experimentally by examining circuit- and neuron-level activity under targeted stimulation using whole-brain calcium imaging and complementary techniques, for example, remains an exciting challenge for the future.

## Supporting information

**S1 Appendix. Additional Tables and Figures**.

## Acknowledgments

We would like to thank Davor Curic for helpful discussions.

## Data Availability

The open *C. elegans* data we used in this study were not collected by us and are described in Refs. [1, 2] and can be obtained from WormAtlas [3]. The computational model used in this study is described in Ref. [30]. Our own code is available here: https://github.com/gourab-sar/C-elegans-synchronization.

## Author contributions

**Conceptualization:** G.K.S., J.D.

**Data Curation:** G.K.S., A.P.

**Formal analysis:** G.K.S., A.P.

**Funding acquisition:** J.D.

**Investigation:** G.K.S., A.P.

**Methodology:** G.K.S., E.T., J.D.

**Project administration:** J.D.

**Resources:** J.D.

**Software:** G.K.S., A.P.

**Supervision:** E.T., J.D.

**Validation:** G.K.S., E.T., J.D.

**Visualization:** G.K.S.

**Writing – original draft:** G.K.S.

**Writing – review & editing:** E.T., J.D.

## Supplementary material

**Table A.** Number of shared neurons between behavioral circuits.

|  | For. loco. | Back. loco. | Chemo. | Mech. | Klino. |
| --- | --- | --- | --- | --- | --- |
| For. loco. | 24 | 0 | 4 | 4 | 2 |
| Back. loco. | 0 | 31 | 10 | 4 | 2 |
| Chemo. | 4 | 10 | 46 | 10 | 13 |
| Mech. | 4 | 4 | 10 | 38 | 0 |
| Klino. | 2 | 2 | 13 | 0 | 23 |

**Table B.** Rich club neurons distribution across FCs for Type I and Type II.

| Type | FC 1 | FC 2 | FC 3 | FC 4 | FC 5 | FC 6 |
| --- | --- | --- | --- | --- | --- | --- |
| Type I | AVAL(R)<br>AVBL(R)<br>DVA<br>PVCL(R) | AIBR<br>RIBL | AVDL(R)<br>AVEL(R) | – | RIAR | – |
| Type II | AVAL(R)<br>PVCL(R) | AIBR<br>RIBL<br>AVBL(R)<br>DVA | AVDL(R)<br>AVEL(R) | – | RIAR | – |

**Table C.** Comparison between the functional communities and the signaling communities.

| Functional Communities | Type | Signaling Communities |  |  |  |  |  |
| --- | --- | --- | --- | --- | --- | --- | --- |
|  |  | SC 1 | SC 2 | SC 3 | SC 4 | SC 5 | SC 6 |
| FC 1 | I (II) | 4 (4) | 2 (1) | 7 (3) | 4 (4) | 0 (0) | 0 (0) |
| FC 2 | I (II) | 8 (7) | 14 (13) | 13 (17) | 6 (5) | 4 (4) | 1 (1) |
| FC 3 | I (II) | 14 (15) | 3 (5) | 7 (7) | 8 (9) | 8 (8) | 0 (0) |
| FC 4 | I & II | 11 | 11 | 6 | 6 | 5 | 1 |
| FC 5 | I & II | 6 | 7 | 0 | 3 | 2 | 0 |
| FC 6 | I & II | 4 | 1 | 1 | 0 | 1 | 0 |

*Note.* The table lists the number of shared neurons between the FCs from both type I and type II with the signaling communities.

**Table D.** Comparison between the signaling communities and the behavioral circuits.

| Behavioral Circuits | Signaling Communities |  |  |  |  |  |
| --- | --- | --- | --- | --- | --- | --- |
|  | SC 1 | SC 2 | SC 3 | SC 4 | SC 5 | SC 6 |
| Forward Locomotion | 1 | 0 | 5 | 2 | 0 | 0 |
| Backward Locomotion | 6 | 2 | 2 | 2 | 0 | 0 |
| Chemosensation | 26 | 5 | 3 | 6 | 0 | 0 |
| Mechanosensation | 9 | 5 | 4 | 1 | 7 | 0 |
| Klinotaxis | 4 | 4 | 6 | 7 | 1 | 0 |

**Fig A.**
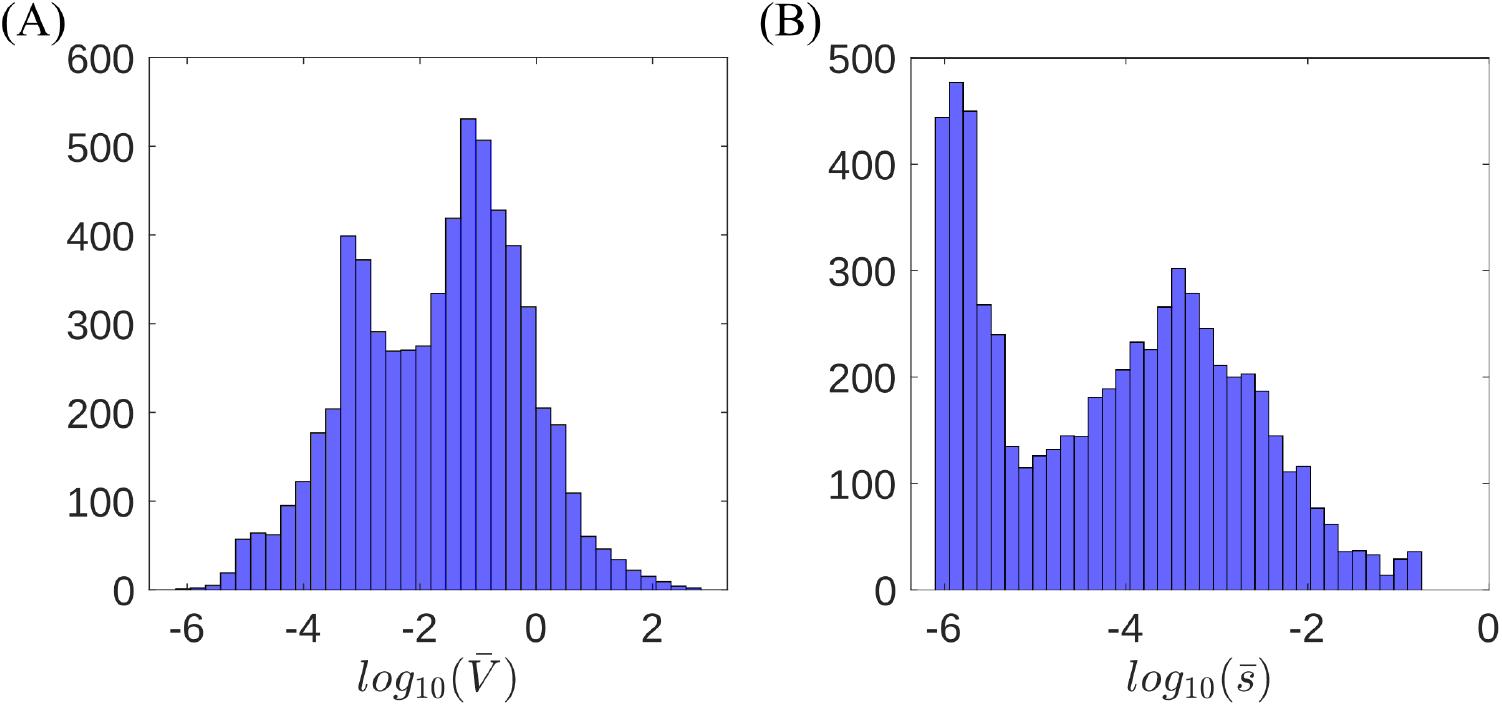
Range of oscillation of the neuron responses. These are calculated as 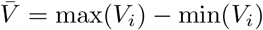 and 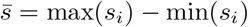 over the last 20% time series data. (A) Histogram of the response voltage of the neuron. (B) Same for the synaptic variable. Data consist of 23 cases for 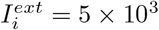where we find oscillatory response.

**Fig B.**
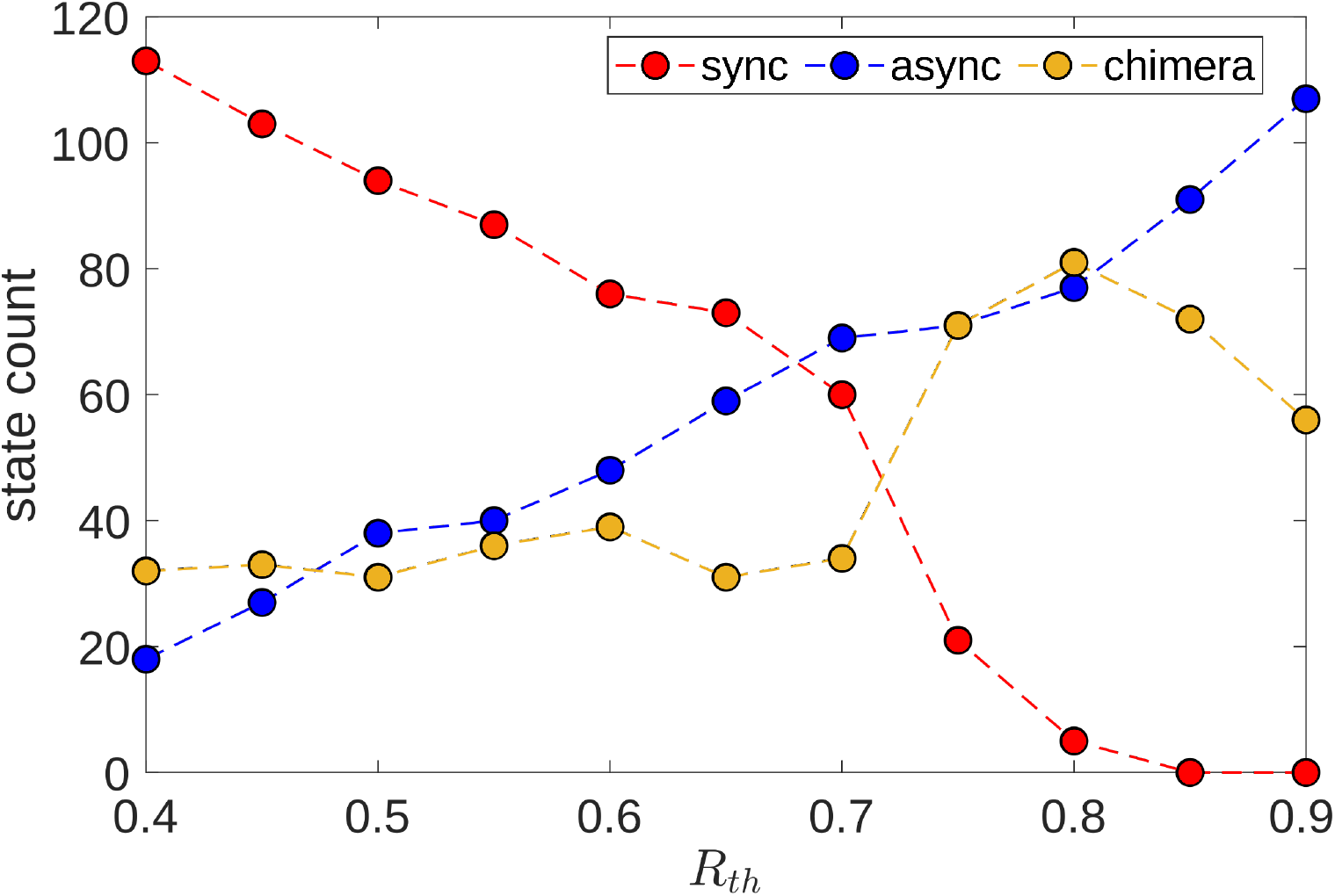
Changes in the count of the sync, async and chimera patterns with *R*_*th*_. Here we vary the synchronization threshold *R*_*th*_ for synchronous community detection using the Louvain algorithm. As *R*_*th*_ is increased, we see the number of sync patterns decrease and async patterns increase which is expected.

**Fig C.**
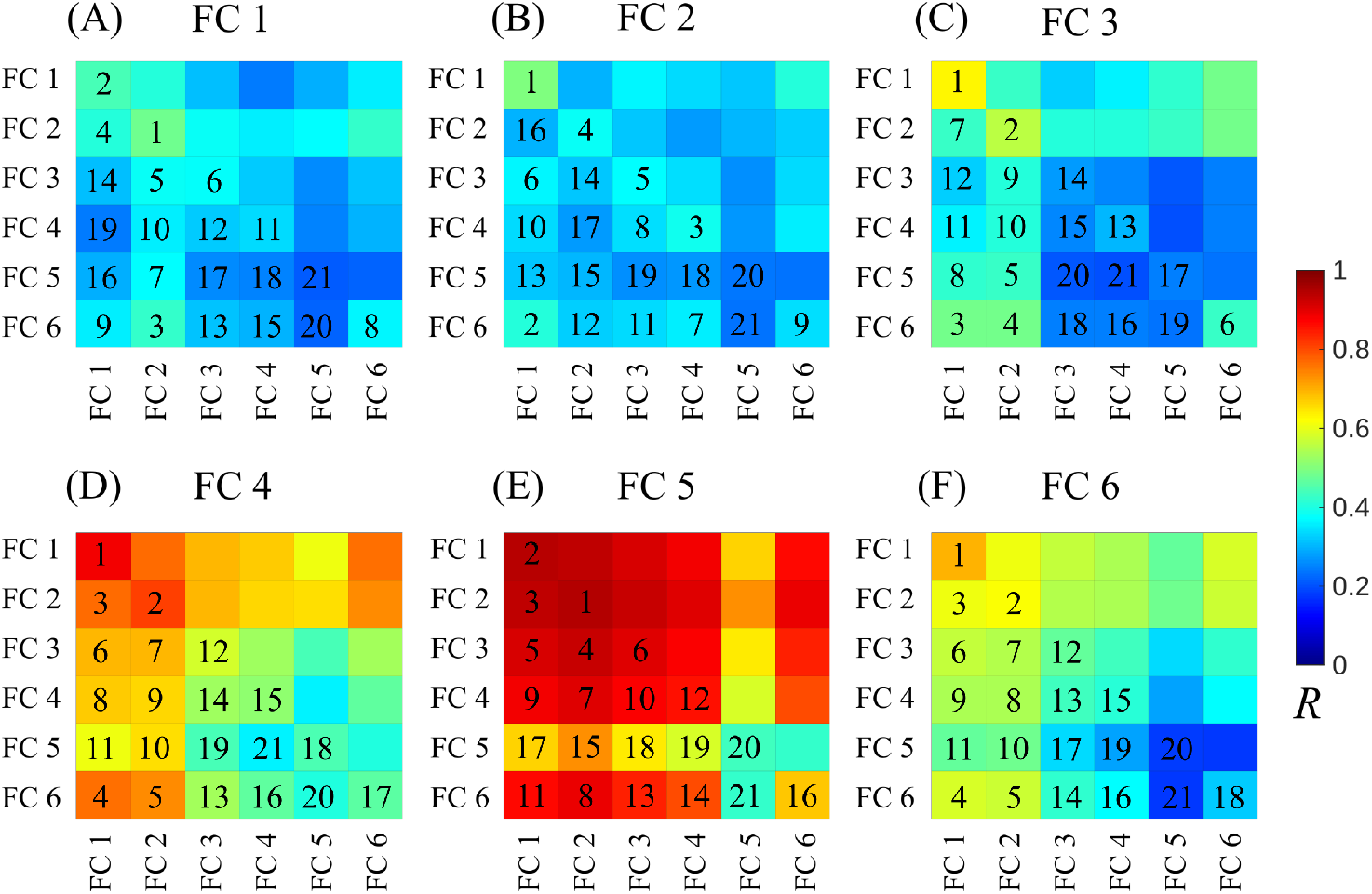
Averaged Kuramoto order matrix *R* for stimulation applied to neurons in different FCs from type II. Panels (A)-(F) correspond to stimulation of neurons in FC 1 through FC 6, respectively. The colorbar indicates the value of *R*, reflecting the degree of synchronization between pairs of FCs. The rank numbers in each matrix are assigned in decreasing order of *R*, where 1 denotes the highest level of synchronization and 21 denotes the lowest.

**Fig D.**
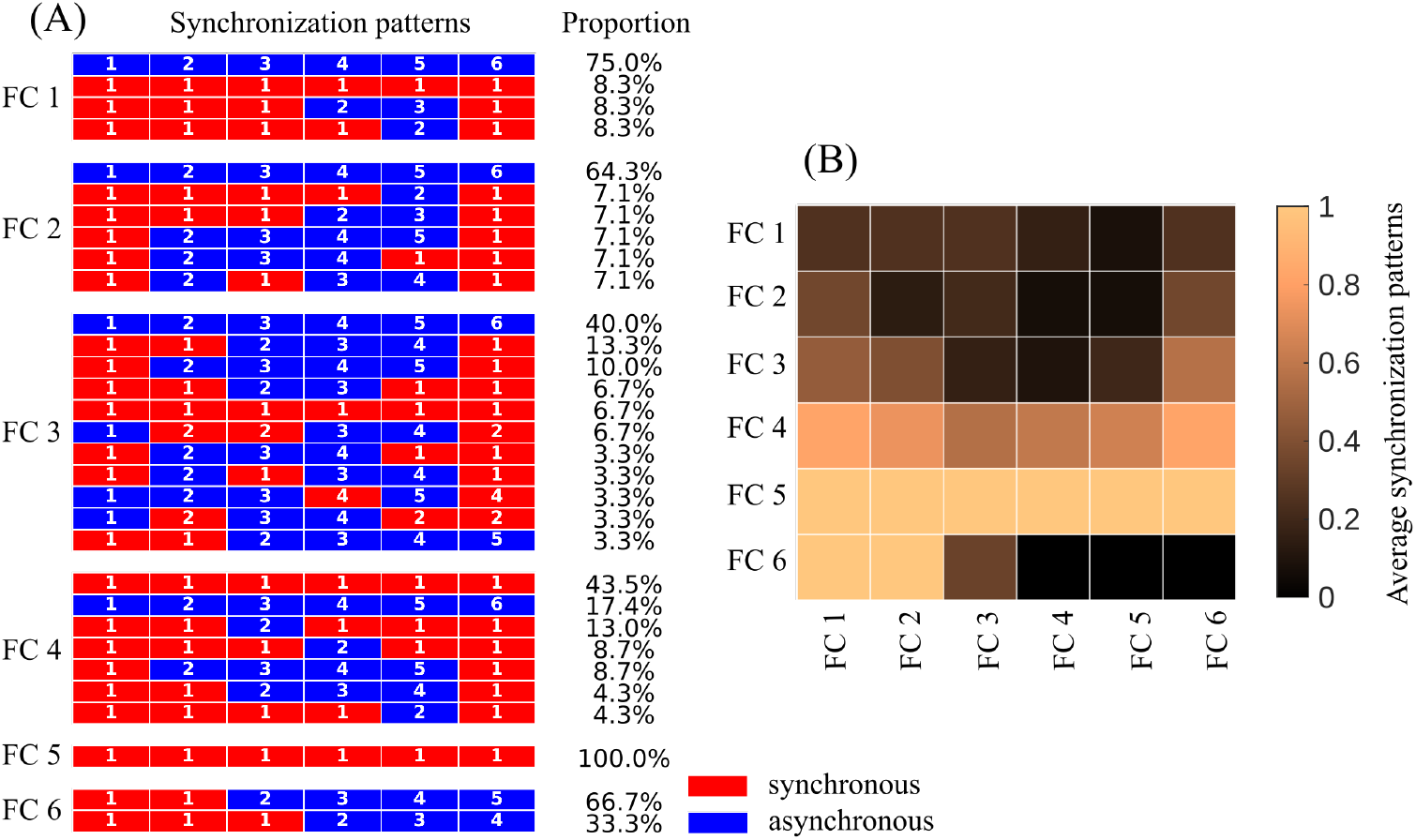
Emergent synchronization patterns following FC-wise stimulation with FCs from WSBM type II. A synchronization threshold *R*_*th*_ = 0.6 is used to generate these patterns. (A) Synchronization pattern types and their proportions for each FC when its neurons are stimulated. (B) Probability of synchronization.

**Fig E.**
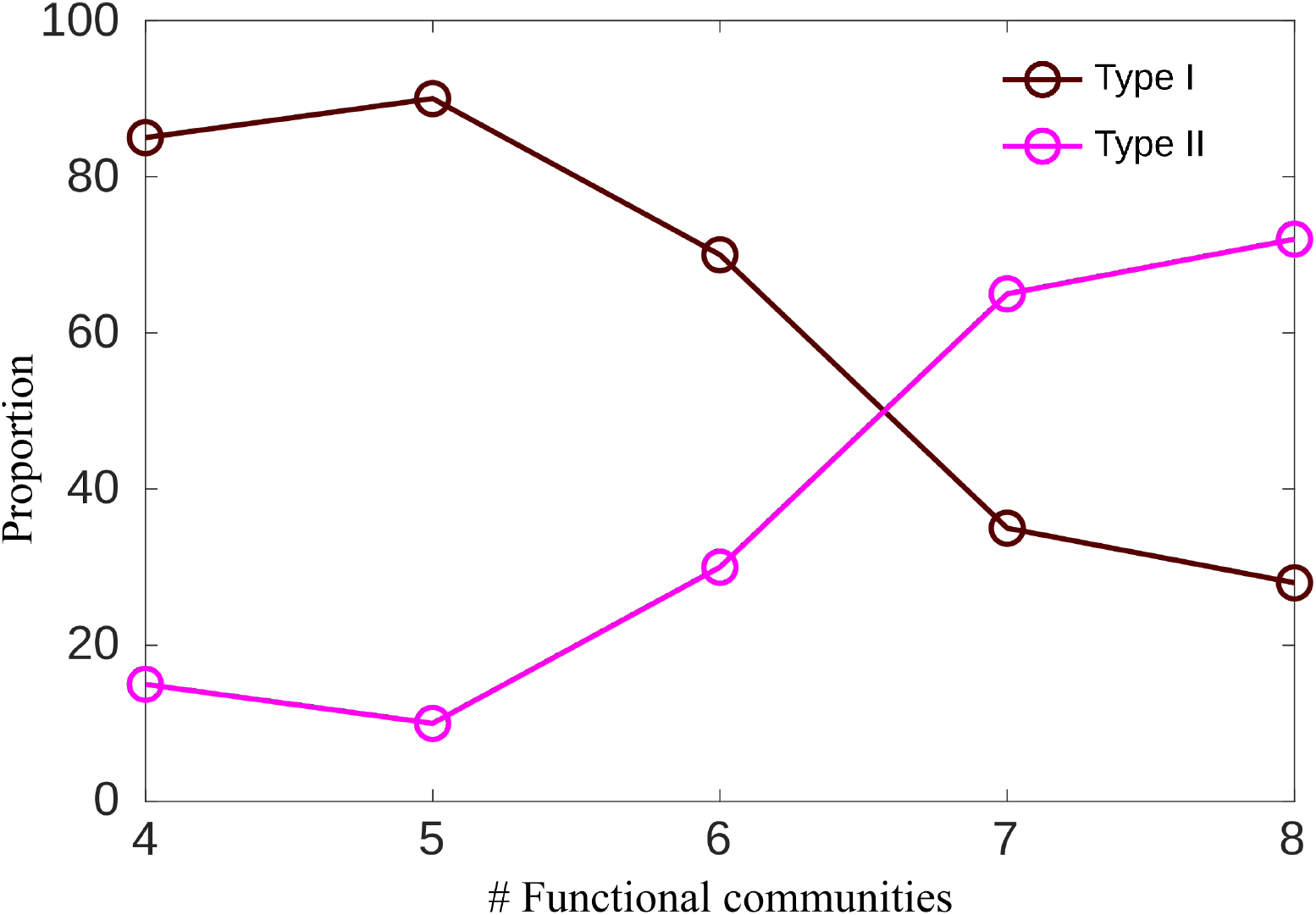
Effect of number of communities. Change in the proportions of Type I and Type II partitions as the number of communities varies.

**Fig F.**
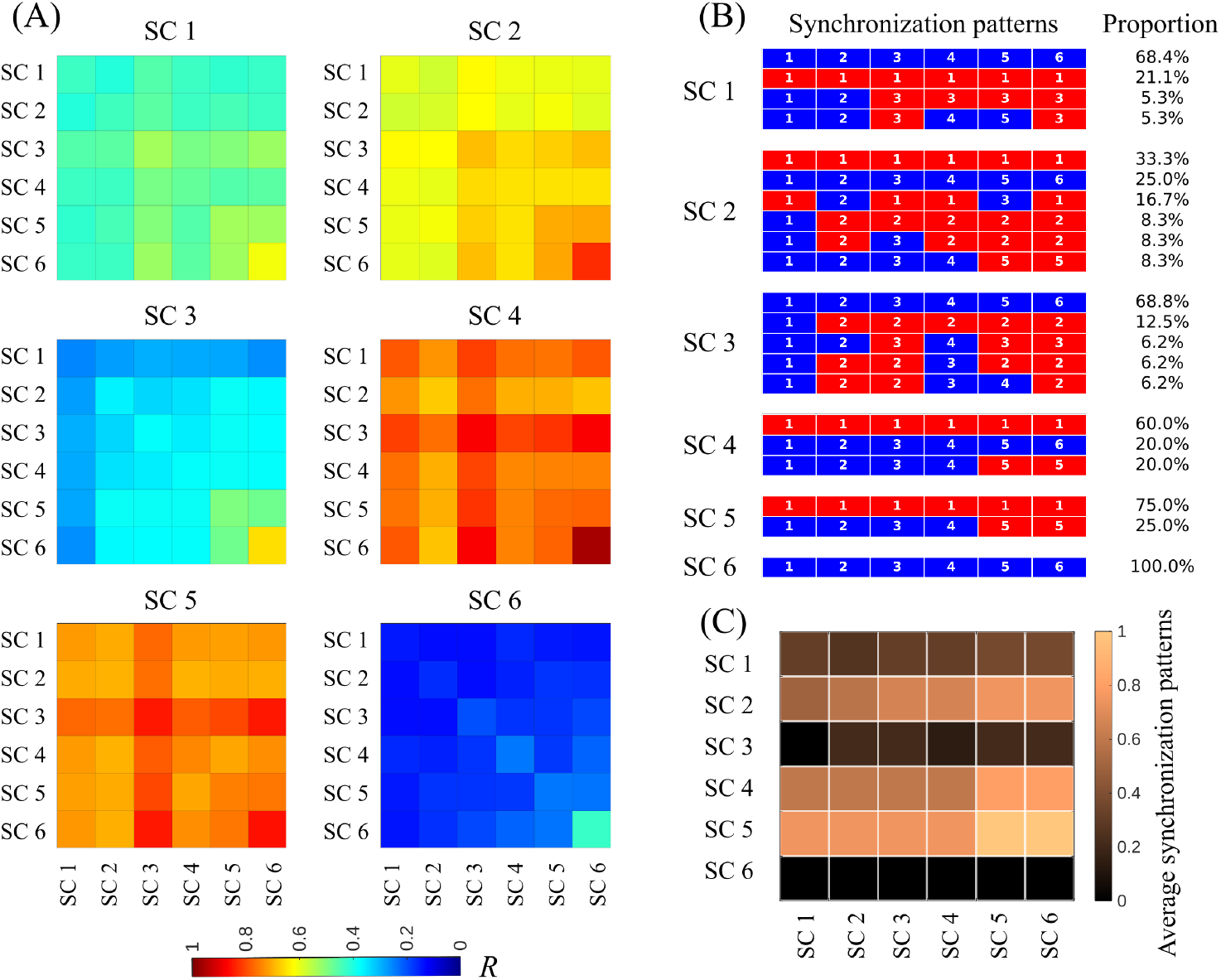
Synchronization patterns in the signaling communities. (A) Average Kuramoto order matrix when neurons of a SC are stimulated. (B) Synchronization patterns and their respective proportions after synchronous community detection using the Louvain algorithm with *R*_*th*_ = 0.6. (C) Probability of a SC being a member of the synchronized group when neurons of some other SC are stimulated. Row-wise driving and column-wise response are shown. The resulting synchronization patterns appear more homogeneous across SCs than what we have observed across FCs.

**Fig G.**
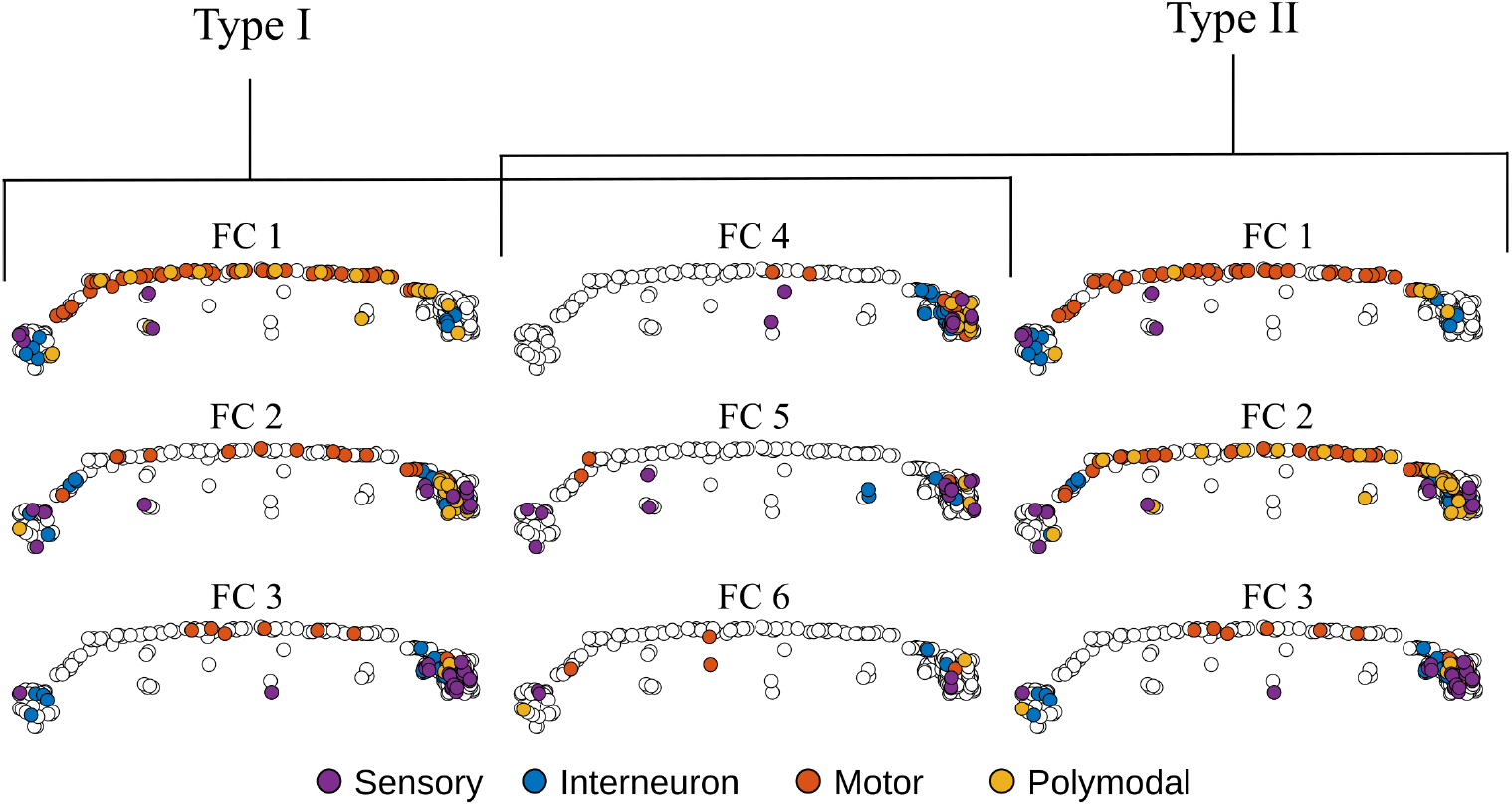
The positional distribution of neurons in each FCs from WSBM type I and type II. The head of *C. elegans* in this figure is towards the right side of the screen from the reader’s point of view.

**Fig H.**
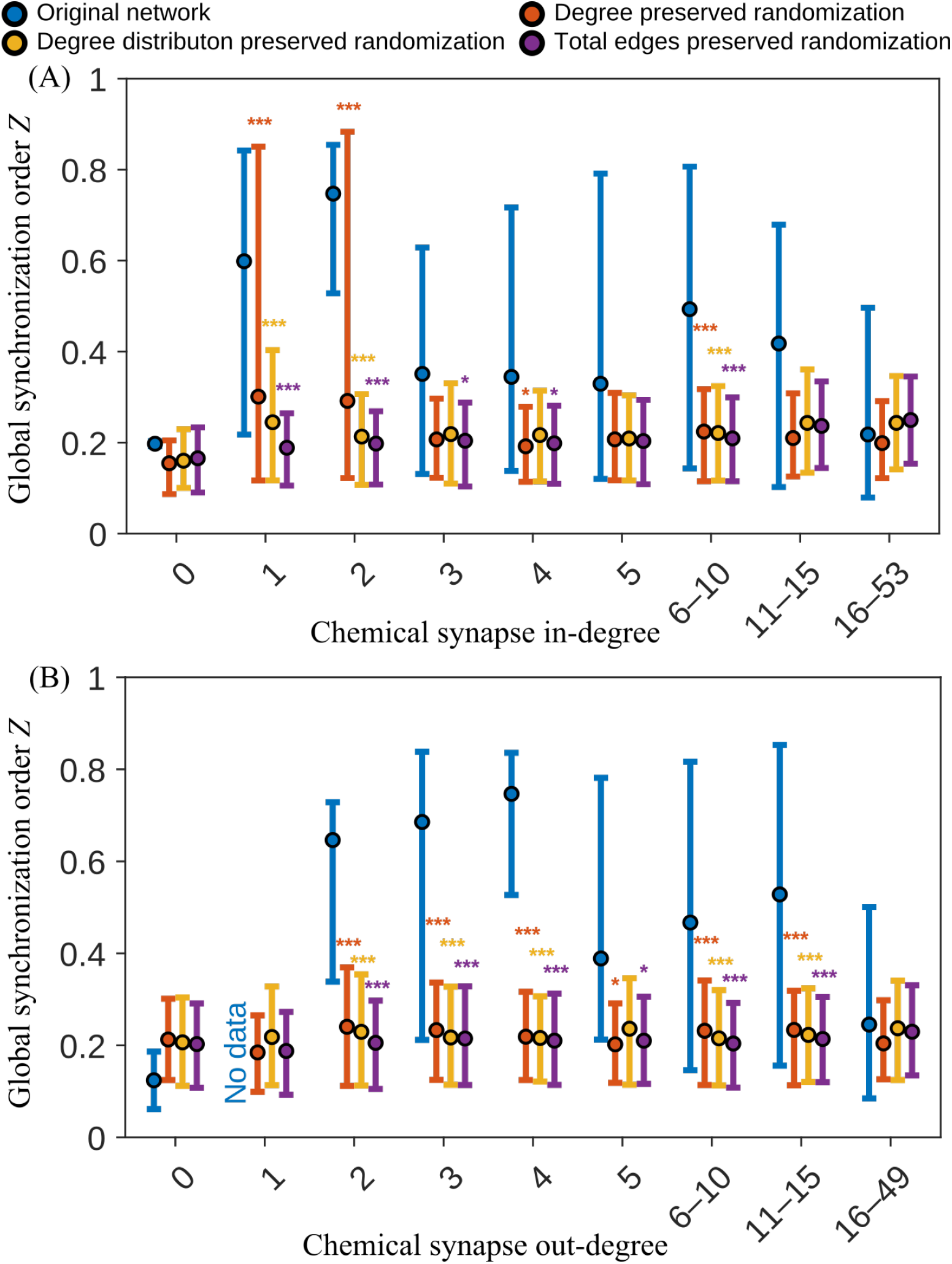
Relation between the global synchronization order *Z* and chemical synapse degree of the stimulated neuron: The original network and the ensemble averages of its three randomized variants are shown, similar to Fig. 7. The dot denotes the mean value and the lower and upper edges represent the 10th and 90th percentiles, respectively. *p* values are used for statistical significance where *, **, and *** denote *p <* 0.05, 0.01, and 0.001, respectively. (A) Chemical synapse in-degree, (B) chemical synapse out-degree.

**Fig I.**
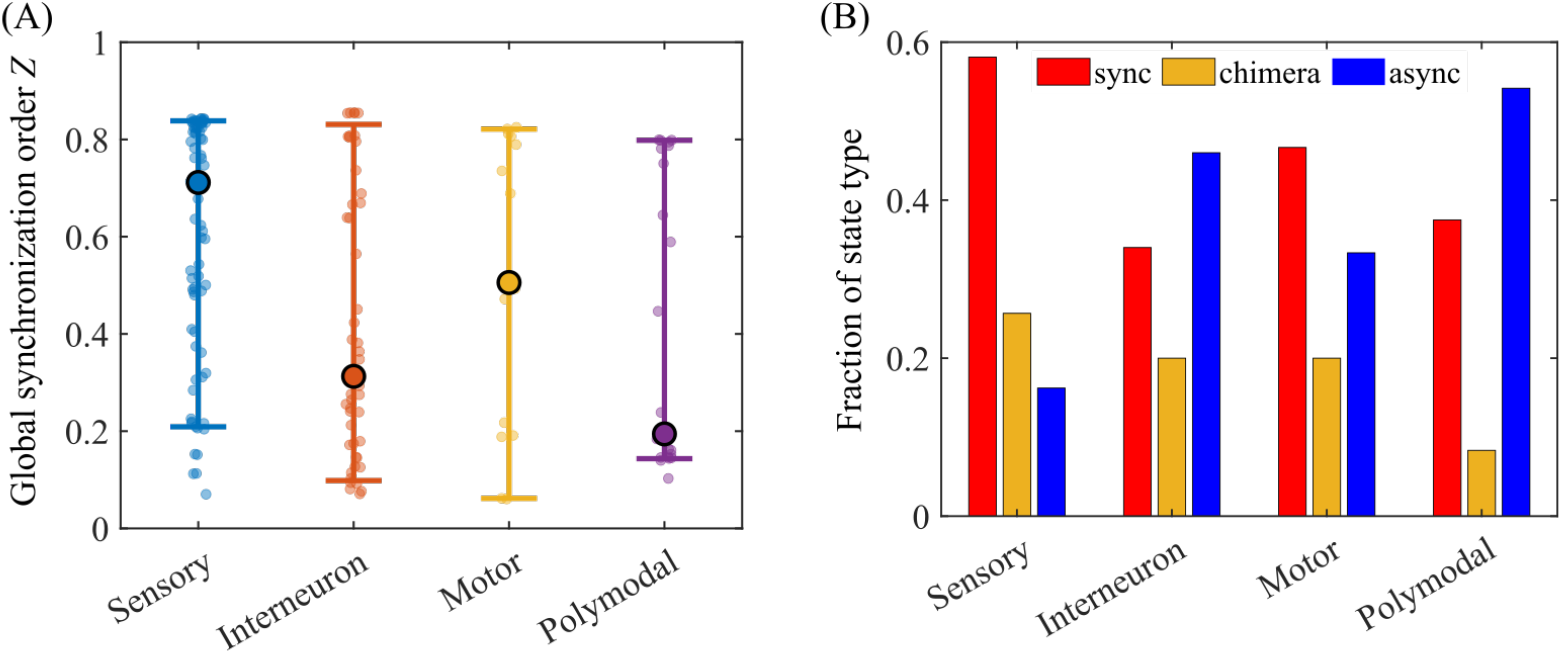
Synchronization patterns and their relation to stimulated neuron types. (A) The global synchronization order *Z* is measured and grouped according to the stimulation of different neuron types. The big dots denote the median values, and the lower and upper edges represent the 10th and 90th percentiles, respectively. (B) The fraction of different synchronization patterns are plotted for each neuron type. A synchronization threshold *R*_*th*_ = 0.6 is used to determine these synchronization pattern using the Louvain synchronous community detection method.

**Fig J.**
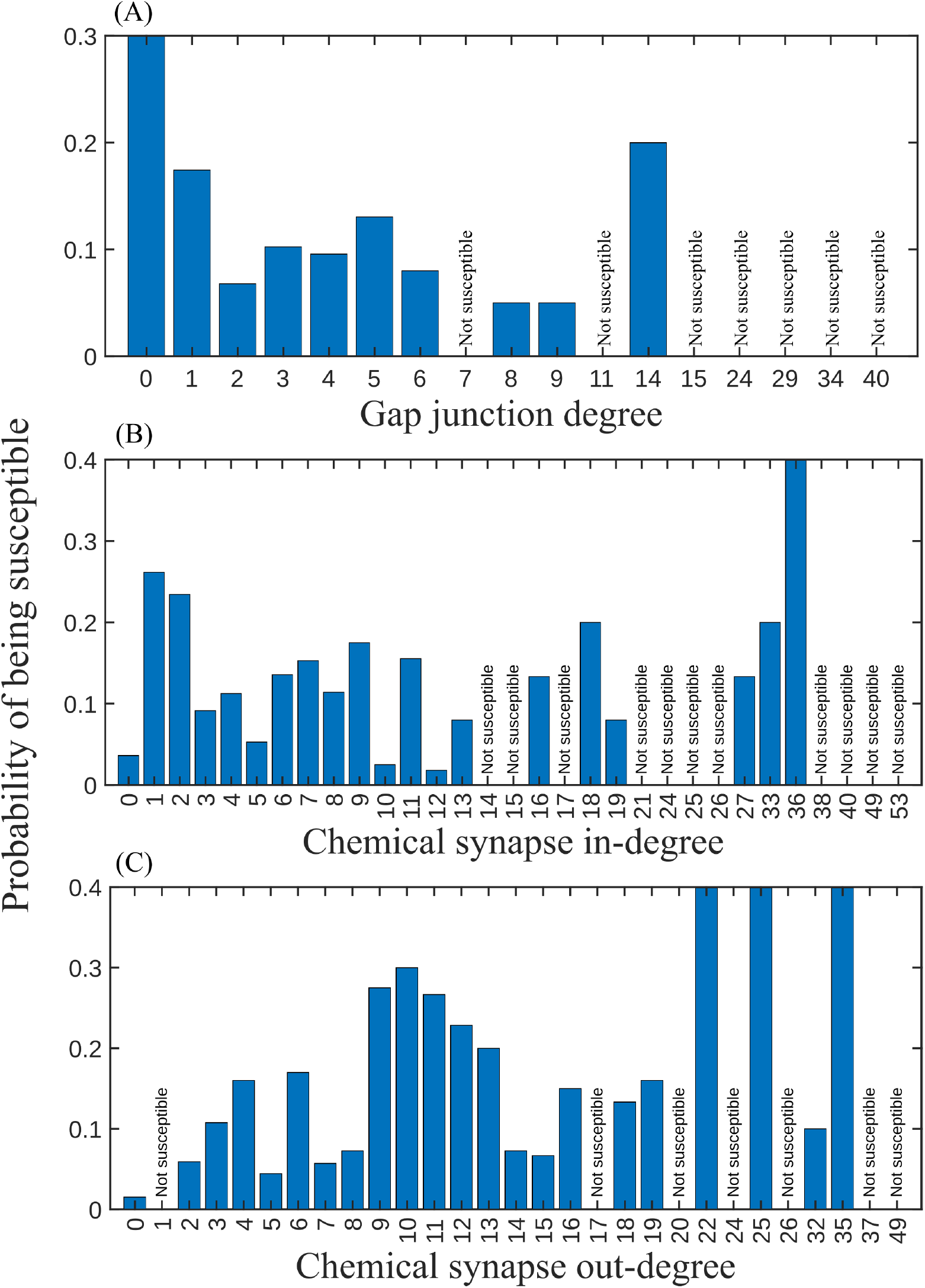
Relation between a neuron being susceptible to stimulation and structural network properties. (A) Low gap junction degree neurons are more probable of being susceptible to stimulation than the high degree ones. There is no such relation of susceptibility with the (B) chemical synapse in-degree, and (C) chemical synapse out-degree.

